# A recombineering pipeline to clone large and complex genes in Chlamydomonas

**DOI:** 10.1101/2020.05.06.080416

**Authors:** Tom Emrich-Mills, Gary Yates, James Barrett, Irina Grouneva, Chun Sing Lau, Charlotte E Walker, Tsz Kam Kwok, John W Davey, Matthew P Johnson, Luke CM Mackinder

**Author notes:** These authors contributed equally to this work (Tom Emrich-Mills, Gary Yates).

## Abstract

The ability to clone genes has driven fundamental advances in cell and molecular biology, enabling researchers to introduce precise mutations, generate fluorescent protein fusions for localization and to confirm genetic causation by mutant complementation. Most gene cloning is PCR or DNA synthesis dependent, which can become costly and technically challenging as genes increase in size and particularly if they contain complex regions. This has been a long-standing challenge for the *Chlamydomonas reinhardtii* research community, with a high percentage of genes containing complex sequence structures, an average genomic GC content of 64% and gene expression requiring regular introns for stable transcription. Here we overcome these challenges via the development of a recombineering pipeline that enables the rapid parallel cloning of genes from a Chlamydomonas BAC collection. We show the method can successfully retrieve large and complex genes that PCR-based methods have previously failed to clone, including genes as large as 23 kilobases, thus making previously technically challenging genes to study now amenable to cloning. We initially applied the pipeline to 12 targets with a 92% cloning success rate. We then developed a high-throughput approach and targeted 191 genes relating to the Chlamydomonas CO_2_ concentrating mechanism (CCM) with an overall cloning success rate of 77% that is independent of gene size. Localization of a subset of CCM targets has confirmed previous mass spectrometry data and identified new pyrenoid components. To expand the functionality of our system, we developed a series of localization vectors that enable complementation of Chlamydomonas Library Project mutants and enable protein tagging with a range of fluorophores. Vectors and detailed protocols are available to facilitate the easy adoption of this method by the Chlamydomonas research community. We envision that this technology will open up new possibilities in algal and plant research and be complementary to the Chlamydomonas mutant library.

## Introduction

The unicellular alga *Chlamydomonas reinhardtii* (hereafter Chlamydomonas) is a widely used model organism for studying photosynthesis, biofuel production, ciliopathies, flagella-powered motility and cell cycle control (Salomé and Merchant, 2019). Its nuclear, chloroplast and mitochondrial genomes are sequenced, well annotated and transformable, and a variety of genetic resources are available to any institution including a close-to-genome-saturating mutant library (Li et al., 2019), extensive -omics based data and a wealth of molecular tools developed over decades by a dedicated research community (Salomé and Merchant, 2019). These collections, data and tools are a vital resource for studies that aim to understand fundamental biological processes, to guide engineering efforts such as improved photosynthetic efficiency and to enable efficient biomolecule production.

Reverse genetic approaches in Chlamydomonas often depend on localizing target proteins to understand spatial distribution and the complementation of mutants to link genotype to phenotype. Both of these methods generally rely on cloning a gene of interest into a plasmid from genomic DNA (gDNA) by PCR, followed by amplification in *Escherichia coli* and reintroduction to Chlamydomonas cells. PCR-based cloning from gDNA presents its own challenges and limitations that are particularly problematic when working with Chlamydomonas nuclear genes, which generally have a high GC content (68% in coding regions), contain one or more introns and can include complex repeating regions (Merchant et al., 2007). On the other hand, cloning from complementary DNA can result in low or no expression of target genes most likely due to lack of introns and lack of regulatory elements (Lumbreras et al., 1998; Schroda, 2019). Some of the challenges associated with PCR-based cloning can be circumvented via gene synthesis, however in many cases the need to include introns, high GC content and gene complexity results in synthesis failure or is prohibitively expensive.

Improved Chlamydomonas target gene and foreign gene (collectively transgenes) expression (e.g., GFP) have been achieved through strain optimization (Neupert et al., 2009), the development of systems with linked transgene and antibiotic resistance gene expression (Rasala et al., 2012; Onishi and Pringle, 2016) and an advanced understanding of transgene silencing (reviewed in Schroda, 2019). Furthermore, release of the Chlamydomonas Golden Gate based Modular Cloning kit has provided a cloning framework and selection of genetic elements to enable labs to rapidly assemble and test transgene constructs (Crozet et al., 2018). Independent of background strain and expression system, it is now clear that inserting or maintaining introns, correct codon usage and promoter sequence are all critical for robust transgene expression (Barahimipour et al., 2015; López-Paz et al., 2017; Baier et al., 2018; Weiner et al., 2018; Schroda, 2019). These considerations have made the cloning of Chlamydomonas target genes directly from gDNA the community standard for mutant complementation and fluorescent protein tagging. However, there are considerable technical hurdles to overcome when working with the expression of large Chlamydomonas genes, predominantly caused by inefficient amplification of gDNA due to gene size, GC content and complexity of target genes (Sahdev et al., 2007). Though modern polymerases have been engineered to overcome sequence challenges (Hommelsheim et al., 2014) they may still suffer from replication slippage events, which are exacerbated by repetitive regions (Levinson and Gutman, 1987; Clarke et al., 2001). In addition to considerations of size, complexity and GC content, cloning native genes based on current genome annotations can be complicated by the abundance of upstream transcription start sites corresponding to possible alternative open reading frames (Cross, 2015) and hence potentially resulting in incorrect target gene cloning.

The results of a recent high-throughput localization study illustrate the challenges of PCR-based cloning of Chlamydomonas nuclear genes (Mackinder et al., 2017). In Mackinder et al. (2017) genes were PCR amplified from start site to stop site using gDNA as the template. Amplicons were then cloned in-frame via Gibson assembly with a fluorescent protein and a constitutive promoter and terminator, resulting in the successful cloning of 298 genes out of an attempted 624 (48% success rate), with most failures at the PCR amplification step. This relatively low success rate led us to develop a cloning platform based on recombination-mediated genetic engineering (recombineering) to enable size and sequence independent cloning of Chlamydomonas genes. Recombineering enables gene cloning by homologous recombination in *E. coli* without PCR amplification of the template, and so is predominantly independent of the target region size. Large-scale recombineering pipelines have been developed for bacterial artificial chromosome (BAC) and fosmid libraries from a broad range of organisms including *Caenorhabditis elegans* (Sarov et al., 2006), *Drosophila melanogaster* (Sarov et al., 2016), human and mice (Poser et al., 2008) and *Arabidopsis thaliana* (Brumos et al., 2020) but are lacking in algae. Our developed pipeline involves making BAC-containing *E. coli* homologous recombination competent by introducing the recombinogenic viral proteins Red α, β and γ from the bacteriophage lambda virus (Yu et al., 2000; Copeland et al., 2001), then retrieving a target sequence via introduction of 50 bp homology regions flanking a linearized plasmid.

We decided to apply our recombineering pipeline to an extended list of putative CO_2_ concentrating mechanism (CCM) genes. The CCM functions to enhance photosynthesis by increasing the concentration of CO_2_ around Rubisco. To achieve this Chlamydomonas actively transports inorganic carbon across the plasma membrane and chloroplast envelope and delivers it as CO_2_ to tightly packed Rubisco within the pyrenoid (Wang et al., 2015). The pyrenoid is a liquid-liquid phase separated compartment (Freeman Rosenzweig et al., 2017; Wunder et al., 2018) essential for CCM function in Chlamydomonas (Meyer et al., 2012; Mackinder et al., 2016). Pyrenoids are widely prevalent in eukaryotic algae and it is thought that approximately 30% of global CO_2_ is fixed within algal pyrenoids (Mackinder et al., 2016). Due to the photosynthetic turbocharging properties of pyrenoid based CCMs there is growing interest in the engineering of them into crop plants to boost yields (Mackinder, 2017; Rae et al., 2017). Recent studies have identified a large number of potential pyrenoid and CCM components (Mackinder et al., 2017; Zhan et al., 2018) that require functional characterization to understand their priority for future synthetic CCM engineering efforts. However, many of these are proving challenging to clone due to size and sequence complexity, making localization and mutant complementation studies difficult.

By applying our pipeline, we have successfully cloned 157 CCM related genes with their native promoters. Cloning appears independent of target gene size and many target genes had multiple complex features that would typically result in PCR failure. The average cloned region was 7.3 kbp and target regions up to 22.7 kbp in size were successfully cloned. The inclusion of the native promoters ensures any upstream open reading frames have been incorporated. The localization of a subset of the proteins encoded by these genes has enabled identification of diverse cellular locations, confirming interaction data (Mackinder et al., 2017) and pyrenoid proteomic data (Mackinder et al., 2016; Zhan et al., 2018). We go on to develop a series of recombineering vectors to enable protein tagging with a range of fluorescent proteins and selection markers for localization, complementation and relative protein abundance studies. The method takes four days to implement, is accessible for any lab equipped for molecular biology and requires no specialized reagents or equipment. The BAC library used in this work and all developed plasmids are available from the Chlamydomonas Resource Centre and a detailed protocol is provided to enable the rapid adoption of this method by research labs to clone nuclear Chlamydomonas genes.

## Results

### Analysis of the Chlamydomonas genome highlights the challenges affecting PCR-based cloning

Cloning Chlamydomonas genes for successful localization and complementation generally requires the amplification of complete open reading frames from gDNA, spanning from their start site to their stop site (ATG-Stop). To gain a better understanding of the challenges involved in cloning Chlamydomonas genes we performed a whole genome analysis of gene size, complexity, intron prevalence, splice variants, and ATG-Stop primer suitability, including comparisons to available datasets and other organisms.

#### Gene size

A major limitation of PCR-based cloning is the target amplicon size. ATG-Stop cloning data from Mackinder et al. (2017) for 624 genes using gDNA as a template and Phusion Hot Start II DNA polymerase (ThermoFisher Scientific) show an association between cloning success and gene size; the average cloned gene was ~2.3 kbp while the average uncloned gene was ~4.5 kbp long (Mann-Whitney *U* = 16059, *P* < 0.001, two-tailed). Extrapolation of PCR efficiency relative to target gene size from Mackinder et al. (2017) to the whole Chlamydomonas genome (version 5.5) indicates that 68% of the genome would be technically challenging to clone via PCR-based methods (Figure 1A), predominantly due to a severe drop off in amplification efficiency for genes >3 kbp long. The largest amplified target in Mackinder et al. (2017) was 8 kbp, and genes at least as large as 9.7 kbp have been cloned before (Kobayashi et al., 2015), but this appears to be highly gene specific. Alternative approaches exist to clone larger genes, such as testing a broad range of PCR conditions and DNA polymerases, amplification in fragments and re-stitching together, cloning from cDNA, and gene synthesis. However, these can be time consuming, costly, have low success rates and may result in no or poor expression.

**Figure 1.**
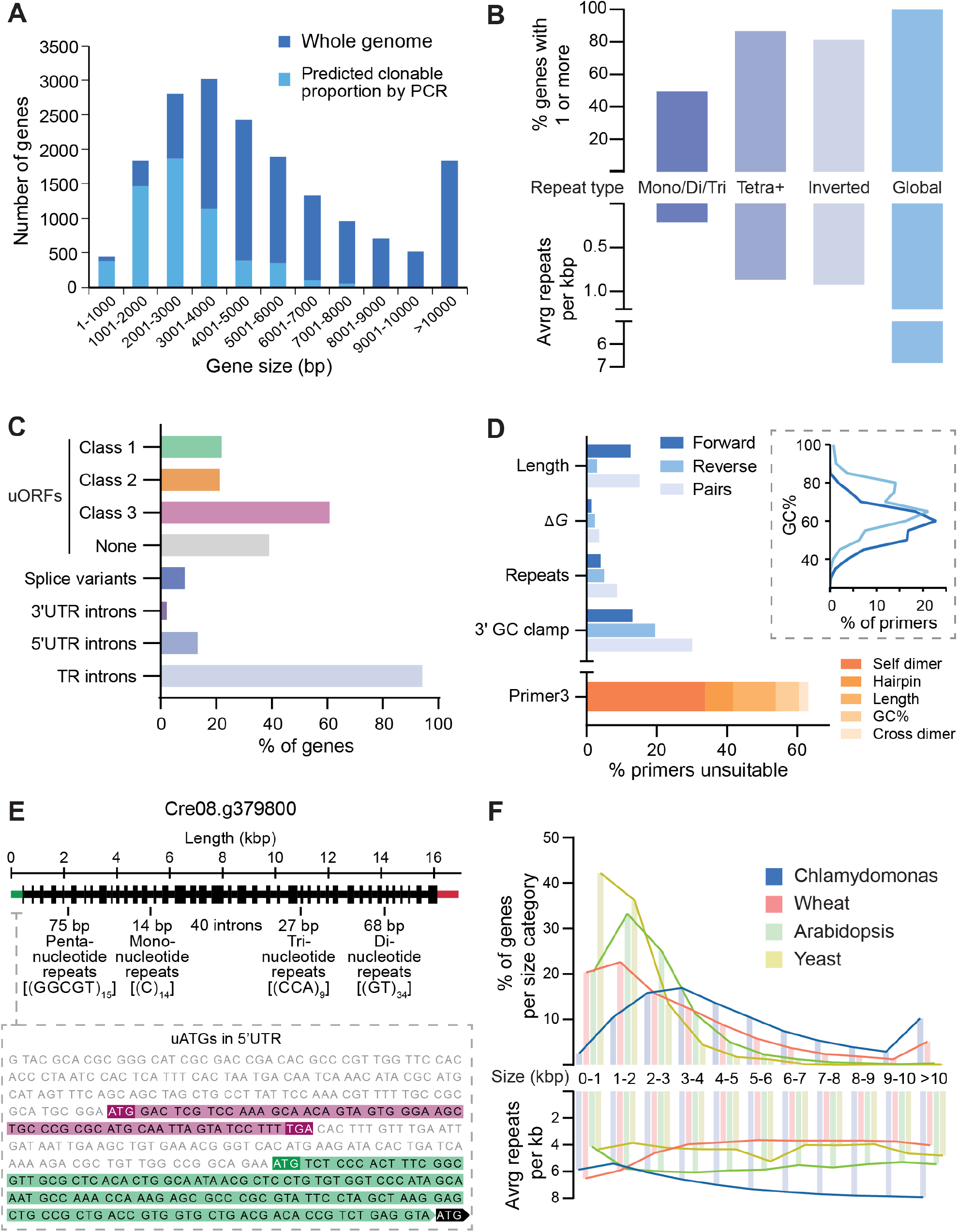
Chlamydomonas nuclear genes are often large, complex, or misannotated, affecting PCR-based cloning attempts and transgene expression success. **A** The distribution of ORF sizes for the 17,741 genes in the Chlamydomonas nuclear genome (dark blue). In each size category, the predicted proportion amenable to cloning (light blue) was estimated by extrapolation of cloning success for 624 CCM-related genes from Mackinder et al. (2017). **B** Genome wide sequence complexity as indicated by the incidence of repetitive sequences in each gene and frequency of repeats per kilobase (kbp). Simple repeats are shown in the left three columns; mono/di/tri refers to tandem repeats with a period length of one, two or three; tetra+ refers to all oligonucleotide tandem repeats with a period length of 4 or more and a total length ≥20 bp. Combining counts for mono-, di-, tri- and tetra+ produces an average value of 1.07 repeats per kbp. Inverted repeats refer to short (20-210 bp) sequences that have the potential to form secondary structures by self-complementary base pairing. Global repeats refer to repetitive sequences masked by the NCBI WindowMasker program (Morgulis et al., 2006), which includes both longer, non-adjacent sequences and shorter, simple repeats (see Methods). Nearly all genes contained detectable repetitive regions using the default WindowMasker settings, with an average of 38.85 per gene. **C** Gene features that complicate correct transgene expression. Top four bars: Potential misannotation of functional start sites shown by percentage of genes containing one or more uORFs of each class (see text). Data is from Cross (2015). Shown below this is the percentage of Chlamydomonas genes with multiple transcript models (splice variants), and those containing introns in the UTRs and translated regions (TR; between start and stop codons). **D** Analysis of a set of ATG-Stop PCR primers designed to clone every gene in the genome from start to stop codon using gDNA as the template (Mackinder et al., 2017). Many primers are predicted to be unsuitable for efficient PCR, as shown by the percentage of forward (dark blue) and reverse (light blue) primers that breach various recommended thresholds associated with good primer design. Pairs (pale blue) are shown for which one or both primers breach the respective thresholds. Thresholds shown pertain to length, secondary structure stability, tandem repeats and 3’ GC content. The inset shows the distribution of GC content of primers in the dataset, illustrating a clear trend in higher GC content at the 3’ end of coding sequences. Below this, the given reason for rejection of primers by the Primer3 check_primers module is shown in orange. Dimer and hairpin values refer to primers rejected for ‘high end complementarity’ and ‘high any complementarity’ errors, respectively. **E** Annotated gene structure of Cre08.g379800. The gene encodes a predicted protein of unknown function but shows examples of several sequence features that contribute to sequence complexity. The unspliced sequence is 16,892 bases long with a GC content of 64.3%. The 41 exons are shown as regions of increased thickness, with 40 introns between them, the annotated 5’UTR in green and the 3’UTR in red. Labels denote selected examples of simple repeats throughout the gene. The inset shows the 5’UTR sequence, displaying examples of two classes of uORFs (see text); class 3 is highlighted in magenta and class 1 in green. For simplicity only one of the seven class 3 uORFs are shown in full. Cre08.g379800 was successfully cloned and tagged using recombineering. **F** A comparison of gene size and complexity between Chlamydomonas, bread wheat (*Triticum aestivum*), *Arabidopsis thaliana* and *Saccharomyces cerevisiae*. Gene sizes were binned as in **A**, and the average number of global repeats per kilobase (kbp) masked by the NCBI WindowMasker program was counted for genes in each size category (Morgulis et al., 2006).

#### Gene complexity

Chlamydomonas genes can be particularly challenging for PCR-based cloning due to high GC content and the presence of numerous repetitive regions. Data from Mackinder et al. (2017) shows that the average GC content for successfully cloned targets by ATG-Stop PCR cloning was 61.4%, while the average for unsuccessful targets was 64.3% - a value exceeded by over 41% of Chlamydomonas nuclear genes. To analyse the genome for repetitive regions, we determined the frequency of simple tandem repeats, inverted repeats, and larger, interspersed repeats. Tandem repeats were assessed by counting individual regions that consist of consecutive mono-, di- or trinucleotide repeats. Mononucleotide repeats shorter than 10 bp and regions of di- and trinucleotide repeats shorter than 20 bp were excluded. Some slight imperfections in the repeating pattern of a region were allowed, with regions that showed ≥90% identity included, such as GGGGGTGGGG. Of the 17,741 coding genes in the genome 8,796 contain one or more mono-, di or trinucleotide repeats (Figure 1B). In terms of prevalence per kilobase, the average Chlamydomonas gene contains 0.21 tandem repeats whereas Arabidopsis contains 0.16 and *Saccharomyces cerevisiae* contains 0.10. Interestingly, if polynucleotide repeats with higher period numbers are counted as well (from tetranucleotide repeats to tandem repeating units of hundreds of base pairs), these values increase 5 fold for Chlamydomonas (1.07 per kbp), 2.5 fold for Arabidopsis (0.39 per kbp) and 3 fold for yeast (0.3 per kbp), highlighting the repetitive nature of the Chlamydomonas genome. Inverted repeats were assessed by counting regions over 10 bp long that are followed closely downstream by their reverse complement, with some mismatches allowed so that regions with ≥90% identity were included. 14,454 genes contain one or more inverted repeats of this kind (Figure 1B), with an average of 0.93 repeats per kbp. To further validate these findings, we analysed the genome for global repeats, which include larger non-adjacent sequences as well as a diverse range of tandem repeats and inverted repeats (Morgulis et al., 2006). With this expanded detection range, Chlamydomonas genes contain an average of 38.9 repeats (6.8 per kbp) whereas Arabidopsis contains 13.7 (5.5 per kbp) and yeast contains 6.0 (4.2 per kbp). Sequence data from Mackinder et al. (2017) for 624 Chlamydomonas genes indicate an association between ATG-Stop PCR cloning success and repeat frequency; the average cloned gene contained 6.1 repeats per kbp whereas the average uncloned gene contained 7.5 repeats per kbp (Mann-Whitney *U* = 24110, *P* < 0.001, two-tailed).

#### Mis-annotation of start sites

Another challenge associated with PCR-based cloning is incorrectly annotated gene models that lead to cloning of a non-biologically relevant sequence. The analysis of transcript models in the Chlamydomonas genome shows that additional ATGs upstream of the annotated start site are highly prevalent (Cross, 2015; Figure 1C top 4 bars). Cross (2015) categorized these potential upstream reading frames (uORFs) into three classes: class 1 uORFs initiate in-frame with the annotated start site, potentially producing an N-terminal extension relative to the annotated gene model; class 2 uORFs initiate out-of-frame with the annotated start site and terminate within the coding sequence; and class 3 uORFs initiate and terminate within the 5’UTR. Data from Cross (2015) on the presence of Kozak sequences preceding class 1 uORFs suggests that approximately half are translated inefficiently *in vivo*. In a PCR-based approach where a constitutive promoter is used, cloning from the wrong ATG may result in an out-of-frame or truncated product, potentially removing essential signal sequences for correct targeting. 57 of the 298 successfully cloned genes from Mackinder et al. (2017) contained a class 1 in-frame ATG upstream of the cloned region, therefore ~10% of cloned regions may have encoded truncated protein products.

#### Introns, UTRs and splice variants

Chlamydomonas genes have a relatively high intron frequency, providing a further challenge for PCR-based cloning. The average gene contains 7.3 introns with an average intron length of 373 bp compared to an average exon length of 190 bp. 94% of genes contain introns between their start and stop codons, 13% of genes contain one or more introns in their 5’UTRs and 3.4% have introns in their 3’UTRs. ATG-Stop cloning would omit introns in UTR regions, potentially missing critical regulatory information. Furthermore, approximately 9% of genes are annotated with two or more transcript models that result from alternative splicing (Figure 1C). This variation would be missed through cloning from cDNA or through gene synthesis that excludes native introns.

#### Unsuitable primers

ATG-Stop PCR cloning of either gDNA or cDNA results in limited flexibility of primer design. Sequence analysis of a set of genome-wide primer pairs for ATG-Stop cloning (Mackinder et al., 2017) indicates that primers are frequently of poor quality and unsuitable for efficient PCR. The average primer in the dataset had a predicted melting temperature (Tm) of 69.2°C and an average GC content of 64.2%. Primer Tm and GC content are expected to be high in comparison to other organisms with less GC-rich genomes, however, many primers also breached recommended thresholds pertaining to length, secondary structure formation, repetitive sequences and 3’ GC content. Primers are shown in Figure 1D (blue bars) as having breached these four thresholds if, (1) they were longer than 30 bp; (2) the free energy (Δ*G*) required to disrupt secondary structure formation (self-dimers, cross-dimers or hairpins) was less than −9 kcal mol^−1^ at PCR-relevant annealing temperatures (66-72°C); (3) they contained mono- or dinucleotide repeats of 5 or more; or (4) their 3’ end contained 5 or more consecutive G/C bases. A stricter set of thresholds is utilized by the Primer3 check_primers module (Rozen and Skaletsky, 2000), which results in the rejection of over 60% of individual primers in the dataset, even when the program is set to ignore predicted annealing temperatures (Figure 1D, orange bar). Under these settings, only 13% of pairs are free from detectable issues in both primers. Interestingly, there is a high GC content mismatch between forward and reverse primers with a considerably higher GC content of reverse primers (Figure 1D, inset).

Many individual genes contain a range of the above features that result in challenges faced during PCR cloning or gene synthesis. Figure 1E shows a gene from chromosome 8 that exhibits several examples and was a target for recombineering. Cre08.g379800 is >16 kbp with 40 introns, contains mono-, di-, tri- and pentanucleotide repeat regions of ≥9 repeats, along with a potential misannotated upstream ATG in the 5’UTR that could initiate a class 1 uORF, as well as seven class 3 uORFs (Cross, 2015).

To further compare whether the challenges faced in Chlamydomonas were similar in other organisms we analysed gene size and gene complexity relative to gene size for the model eukaryote *S. cerevisiae*, the model plant Arabidopsis and the ~17 Gb hexaploid genome of *Triticum aestivum* (bread wheat). Figure 1F shows that Chlamydomonas has a higher proportion of long genes and fewer short genes than the three other genomes tested, along with a considerably higher average gene size for Chlamydomonas (5322 bp versus 1430 bp for yeast, 2187 bp for Arabidopsis and 3521 bp for chromosome-assigned genes in wheat). Unlike wheat, Arabidopsis and yeast, Chlamydomonas genes show a trend of increasing complexity per kilobase for longer genes (Figure 1F), potentially in line with an increase in average UTR length as gene length increases (Salomé and Merchant, 2019).

### Recombineering pipeline development

To overcome the challenges associated with PCR-based cloning we developed a high-throughput recombineering pipeline for large-scale parallel cloning of Chlamydomonas nuclear genes from BACs with their native promoter regions intact. During pipeline development we decided to pursue a simplified 1-step DNA retrieval recombineering approach rather than a BAC editing approach (i.e. Poser et al., 2008; Brumos et al., 2020) for several reasons: (1) Using a gene retrieval method enables all cloning to be performed in the BAC host *E. coli* strain, thereby avoiding BAC purification, which can be timely and low yielding; (2) assembled constructs contain only the gene of interest making them considerably smaller than the original BAC, this allows a medium copy origin of replication to be used that improves ease of handling, and the smaller constructs minimize DNA fragmentation during Chlamydomonas transformation (Zhang et al., 2014); (3) BACs contain many genes, with additional copies of adjacent genes to the gene of interest potentially having an unwanted phenotypic impact on transformed Chlamydomonas lines; (4) the backbone of the available BAC collection lacks a suitable Chlamydomonas selection marker, therefore additional BAC editing to insert a suitable selection marker (Aksoy and Forest, 2019) or inefficient and poorly understood plasmid co-transformation strategies would be required for selection; and (5) a typical BAC engineering approach would require two recombination steps, which would increase pipeline time, decrease pipeline efficiency and add further challenges due to the repetitive nature of the Chlamydomonas genome.

The simplicity of our pipeline enables completion in four days using only generic reagents. The final recombineered construct is a vector containing the target region (typically including the native promoter and open reading frame) recombined in-frame with a downstream fluorescent protein followed by the *PSAD* terminator (see Figure 2 for a pipeline schematic and Supplemental Method 1 for a detailed protocol). Our pipeline has four key steps: (1) *E. coli* harbouring a BAC containing the gene of interest is made recombination competent by transformation with the pRed vector containing the lambda viral *exo*, *beta* and *gam* genes (Redαβγ) and *recA* (Sarov et al., 2006) (Figure 2A); (2) Redαβγ and *recA* induction by arabinose followed by transformation with a linear tagging cassette including 50 bp homology arms to the target gene (Figure 2B); (3) kanamycin selection for successful recombination events and temperature inhibition of the pRed pSC101 replication origin to minimise further undesired recombination (Figure 2C); and (4) plasmid isolation and verification via restriction digest and junction sequencing (Figure 2D).

**Figure 2.**
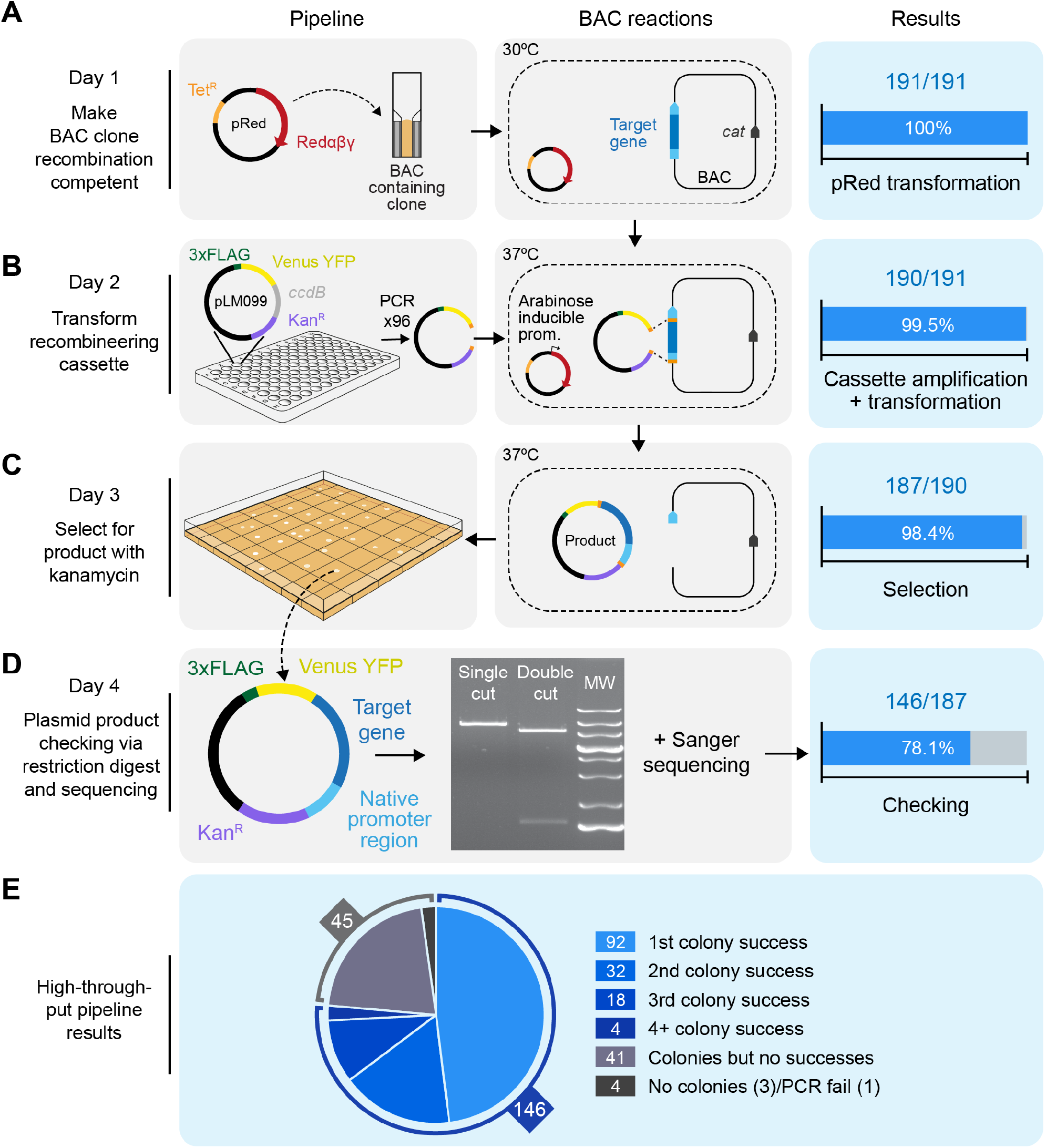
We developed a high throughput recombineering pipeline for generating Venus-tagged fusion proteins with native promoter regions intact. **A** On day 1, BAC clones containing target genes are made recombineering competent by transformation with the pRed plasmid, which encodes the viral recombinogenic Redαβγ genes and *recA* under the control of an arabinose inducible promoter. Transformation efficiency shown on the right-hand side relates to BAC clones that yielded colonies after selection with tetracycline and chloramphenicol. *Cat*: the chloramphenicol resistance gene in the backbone of every BAC clone in the BAC library. **B** On or before day 2, the recombineering cassette is amplified from pLM099 using primers that contain 50 bp homology arms complementary to regions flanking the target gene (shown in orange); one >2000 bp upstream of the annotated ATG and one at the 3’ end of the coding sequence. On day 2, BAC-containing cells are electrotransformed with the recombineering cassette after induction with L-arabinose. Recombination between the BAC and the cassette results in a plasmid product containing the target gene in frame with CrVenus-3xFLAG and under its native promoter. Efficiency shown at this stage relates to PCR reactions that yielded efficient amplification of the recombineering cassette. **C** On day 3, colonies containing plasmid products are isolated. Efficiency at this stage relates to the number of transformations that yielded colonies after selection with kanamycin. **D** On day 4, plasmid products are extracted from cells, screened by enzymatic digestion and confirmed by sequencing. Efficiency shown at this stage relates to correct digest patterns with single and double cutting restriction enzymes. MW: molecular weight marker. **E** Overall efficiency split into number of colonies screened via restriction digest. For 74% of target regions, the correct digest pattern was observed from plasmids isolated from the first, second or third colony picked per target. For 3% of targets, analysing >3 colonies yielded the correct product.

The original tagging cassette consists of the codon optimized YFP CrVenus, a 3xFLAG tag, the *PSAD* terminator, the paromomycin selection marker (*AphVIII*), the p15A medium-copy-number origin of replication and the kanamycin resistance gene (*Kan^R^*). Amplification of the tagging cassette from pLM099 is performed using primers containing 50 bp homology arms corresponding to regions flanking the target gene; one designed at least 2,000 bp upstream of the start codon to encompass the native 5’ promoter and UTR region, and one at the 3’ end of the coding region (immediately upstream of the stop codon). To minimise false positives due to pLM099 carryover, pLM099 contains the *ccdB* counter selection gene (Bernard and Couturier, 1992). In addition, the cassette includes an I-SceI restriction site. I-SceI has an 18 bp recognition site not found within the reference Chlamydomonas genome (strain CC-503) and allows cassette linearization prior to transformation into Chlamydomonas (see Figure S1 for a detailed map of pLM099).

We initially tested our pipeline on 12 targets. To ensure that the BAC library (available from the Chlamydomonas resource centre; https://www.chlamycollection.org/) was correctly mapped we performed PCR to check for the presence of the 5’ and 3’ ends of our target genes (Figure S2A). We next implemented the pipeline according to a small-scale batch protocol (Supplemental Method 1A). For all targets except one, most picked colonies gave a correct banding pattern after restriction digest (Figure S2B). After sequence confirmation we successfully cloned 11 out of our 12 targets, resulting in a 92% success rate (Figure S2C). Confident that our recombineering method was robust we pursued the development of a large-scale pipeline that would allow the parallel tagging of genes with most steps achievable in 96-well format.

### Successful large-scale application of the recombineering pipeline

To test the efficiency of the pipeline we shortlisted 191 genes which could be mapped to a clone from the Chlamydomonas BAC library. To more easily identify BACs within the library that contain a target gene we designed a Python script (BACSearcher; Supplemental Code) and have outputted the five smallest BACs for all targets in the genome in Supplemental Data Set 1, revealing that 86% of nuclear genes are covered by at least one BAC (87% if BACs are included that terminate within 3’UTRs). BACSearcher also enables automated design of primers containing 50 bp homology regions to target genes in optimal positions; the script reports suitable 5’ homology regions 2000-3000 bp upstream of the annotated start codon and takes into account local DNA complexity features, including mono- and dinucleotide repeating runs and GC content. This feature can be easily modified to design 5’ homology regions further upstream of the target (see Supplemental Method 2A) The length of 50 bp is short enough to design into an oligonucleotide but long enough to be unlikely to share homology with more than one site within a BAC. Supplemental Data Set 1 includes sequences for the top five optimal 5’ homology regions for each target, all >2000 bp upstream of the start codon, along with the corresponding 50 bp 3’ homology region. In addition, four pairs of primer sequences are included that can be used to check for the presence of each target in a BAC.

Our 191 targets were primarily chosen based on our 2017 association study for CCM components (Mackinder et al., 2017), transcriptomics (Brueggeman et al., 2012; Fang et al., 2012) and pyrenoid proteomics (Mackinder et al., 2016; Zhan et al., 2018). *E. coli* strains containing the correct BAC as identified by BACSearcher were recovered from the BAC library and processed in parallel using 96-format culturing plates. To optimise the efficiency of our high-throughput pipeline, we successively ran the pipeline three times removing successful targets once confirmed. Supplemental Method 1B provides a detailed protocol for the optimized high-throughput pipeline. In summary, 100% of our 191 target BAC lines were made recombination competent (Figure 2A), and out of the 191 target genes, one gene-specific tagging cassette failed to amplify (Figure 2B), likely due to the formation of secondary structure(s) within the 50 bp homology regions of the primers. Of the 190 that amplified successfully, 187 yielded colonies after selection with kanamycin (Figure 2C). Validation by enzymatic digestion confirmed that 146 of these lines contained correct recombineering plasmid products. For the 146 correctly recombineered lines, picking just a single colony gave a 63% success rate, screening a second colony increased the success rate to 85% and a third colony gave a 97% success rate, for a small proportion of targets screening >3 colonies led to the identification of a correctly recombined construct (Figure 2E). Recombineering plasmid products from the 146 successful lines were extracted and their junctions confirmed by Sanger sequencing. Our high-throughput pipeline had an overall efficiency of 76%, an average recombineered region of 7259 bp and a maximum cloned length of 22,773 bp corresponding to gene Cre10.g427850 (Supplemental Data Set 2).

During pipeline development, we found that optimising bacterial growth prior to transformation with the recombineering cassette was critical (see protocol notes in Supplemental Method 1). Also, in 14 cases, using an alternative BAC from the library yielded success. Restriction digest analysis of plasmids isolated from incorrectly assembled recombineering events suggested that cloning could fail due to a broad range of reasons including cassette recircularization, cassette duplication, cassette insertion into the BAC or retrieval of incorrect target regions. Increasing homology arm length, using alternative homology arms and using alternative BACs are potential solutions to overcome incorrect recombineering for specific targets. Supplemental Data Set 1 provides up to five options for homology arms and up to five available BACs per gene and can be easily modified to increase homology arm length (see Supplemental Method 2A).

### Cloning success is size independent and tolerant of sequence complexity

To investigate if our developed recombineering approach was gene size and complexity independent, we compared our successful targets against unsuccessful targets (Figure 3). The results show that there is no significant difference in the region lengths between cloned and uncloned regions (Figure 3A; Mann-Whitney *U* = 3306, *P* = 0.38, two-tailed), indicating that our method is target size independent. This contrasts to the clear effect of target size on cloning success for our previous PCR-based cloning data (Figure 3A; Mackinder et al., 2017). To explore if our recombineering method was affected by gene complexity we compared our cloning success to the number of simple and global repeats per kilobase in target genes. Our method appears far more tolerant of repetitive sequences than PCR-based cloning, both in the per-kilobase prevalence of simple and global repeats and in the number of repeats per target region (Figure 3B and C). For our recombineering pipeline there is no significant difference detectable in the average repeat prevalence per kilobase between cloned and uncloned regions (Mann-Whitney *U = 2899, P* = 0.19, twotailed), while there is a clear negative effect on PCR-based cloning success for targets with over ~4.8 repeats per kbp (Figure 3B). For the most repetitive targets we analysed (>9 repeats per kbp), our recombineering cloning efficiency stayed above 60%; an efficiency over three times higher than PCR-based cloning. Extrapolation of these data, overlaid with the genome wide distribution of repeat frequencies, indicates that a large proportion of genes technically challenging for PCR-based cloning due to high repeat frequencies can be cloned by recombineering (Figure 3B).

**Figure 3.**
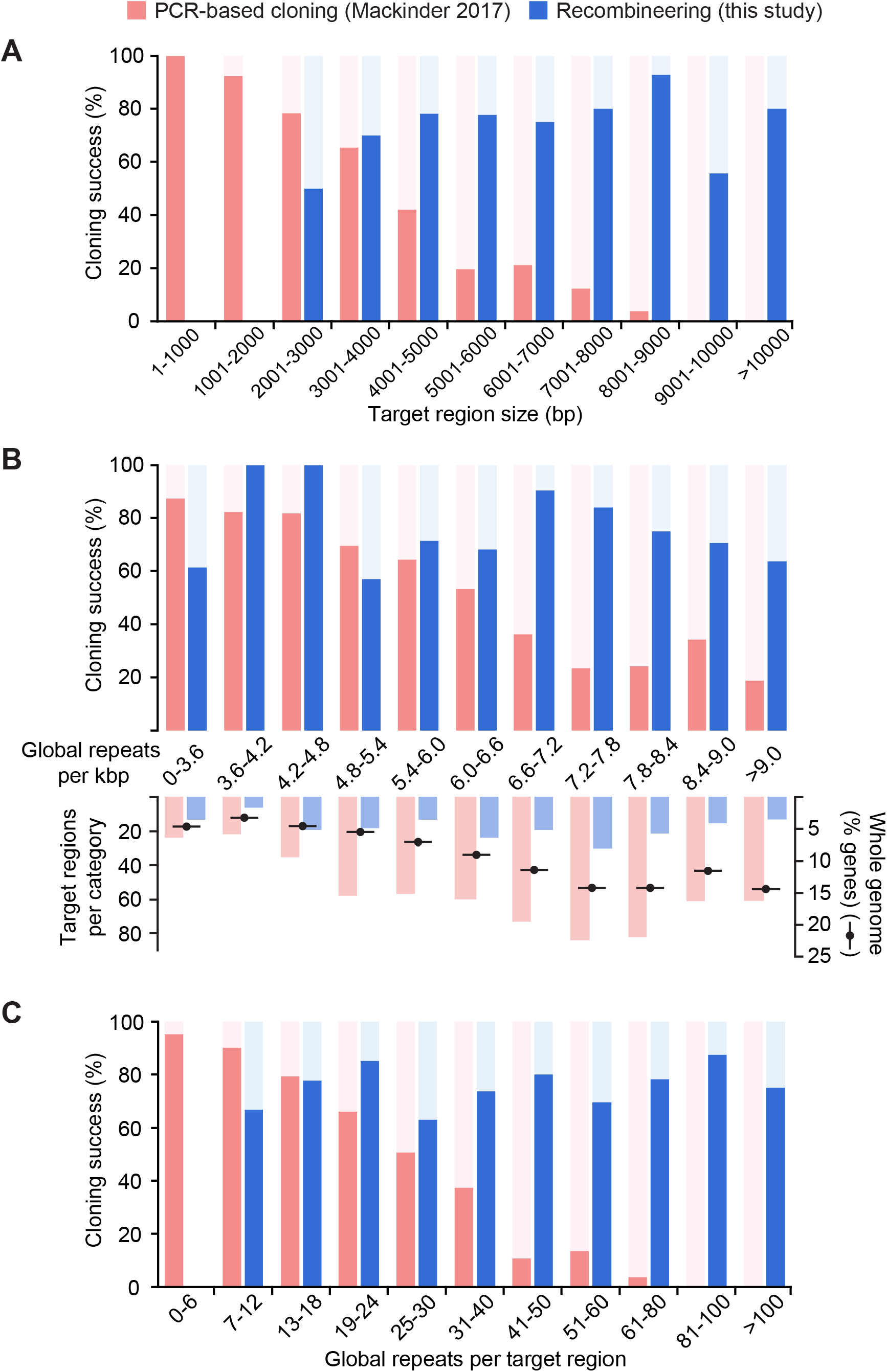
Our recombineering pipeline is target gene size independent and tolerant of sequence complexity **A** The size distribution of successfully PCR-cloned coding sequences (Mackinder et al., 2017; red) or recombineered regions (this study; blue) are shown. Regions cloned by recombineering include ~2 kbp of flanking DNA upstream of the annotated start codon to incorporate native 5’ promoter sequences. A severe drop in PCR-based cloning efficiency can be seen for templates >3 kbp long, whereas recombineering cloning efficiency does not show size dependency. No recombineering target regions were less than 2000 bp long due to inclusion of native 5’ promotor sequences. **B** As above but showing the dependence of cloning success on the per-kilobase frequency of repeats masked by the NCBI WindowMasker program with default settings (Morgulis et al., 2006). The number of target regions per repeat category is shown beneath this, overlaid with the percentage of Chlamydomonas genes in each category. The distribution of targets for this study and our previous PCR-based cloning attempt (Mackinder et al., 2017) gives a reasonably close representation of the whole genome distribution. Almost a third of nuclear genes contain 7.2-8.4 repeats per kbp; this peak corresponds to a clear drop in PCR-based cloning efficiency, but to a high recombineering efficiency of 75-85%. Data for repeats per kbp was continuous and there are no values present in more than one category. **C** As above but showing the number of simple and global repeats masked by WindowMasker per template. Data are binned to provide a higher resolution for the lower value categories, since the targets for PCR-based cloning were enriched in targets with low numbers of repeats. As in **A**, a severe negative trend in PCR-based cloning efficiency can be seen, reflecting a strong positive correlation between repeat number and region size. No negative association is present for recombineering cloning efficiency, likely illustrating the benefit of avoiding size- and complexity-associated polymerase limitations. No recombineering target regions contained fewer than 6 repeats.

### Localization of Venus-tagged proteins

To assess the validity of the pipeline for localization studies we transformed wild type Chlamydomonas cells with a subset of linearized recombineering plasmid products tagged at the C-terminus with CrVenus (Figure 4A). Paromomycin resistant colonies were directly screened for YFP fluorescence on transformation plates, picked, grown in TP media at air-levels of CO_2_ (~0.04%), and then imaged by fluorescence microscopy to examine the localization pattern (Figure 4B). Transformed genes were selected based on previous affinity purification mass spectrometry data (Mackinder et al., 2017) and pyrenoid proteomics data (Mackinder et al., 2016; Zhan et al., 2018). The localization data supports the proteomics data with PSAF (Photosystem I subunit F; Cre09.g412100), ISA1 (Isoamylase 1; Cre03.g155001) and Cre10.g435800 present in the pyrenoid. PSAF is a core transmembrane subunit of photosystem I. As expected PSAF shows strong colocalization with chlorophyll outside of the pyrenoid, however in addition it clearly localizes to the thylakoid tubules traversing the pyrenoid. Interestingly, in the pyrenoid tubules the chlorophyll signal is minimal, particularly at the “pyrenoid tubule knot” where the tubules converge (Engel et al., 2015). These data along with the localization of other PSI and PSII components to the pyrenoid tubules (Mackinder et al., 2017) suggest that the tubules contain both PSI and PSII but that chlorophyll-containing light harvesting complexes found within the pyrenoid may be quenched or at low abundance. Tagged Cre17.g702500 (TAB2), a protein linked to early PSI assembly (Dauvillée et al., 2003) and which was identified as an interactor with PSBP4 found within and at the periphery of the pyrenoid (Mackinder et al., 2017), was also enriched at the pyrenoid. Interestingly, the location of TAB2 is not just restricted to the pyrenoid periphery but is also found within the pyrenoid forming distinct small foci (Figure 4B). This may indicate that early PSI assembly could be occurring within the pyrenoid as well as at the pyrenoid periphery (Uniacke and Zerges, 2009).

**Figure 4.**
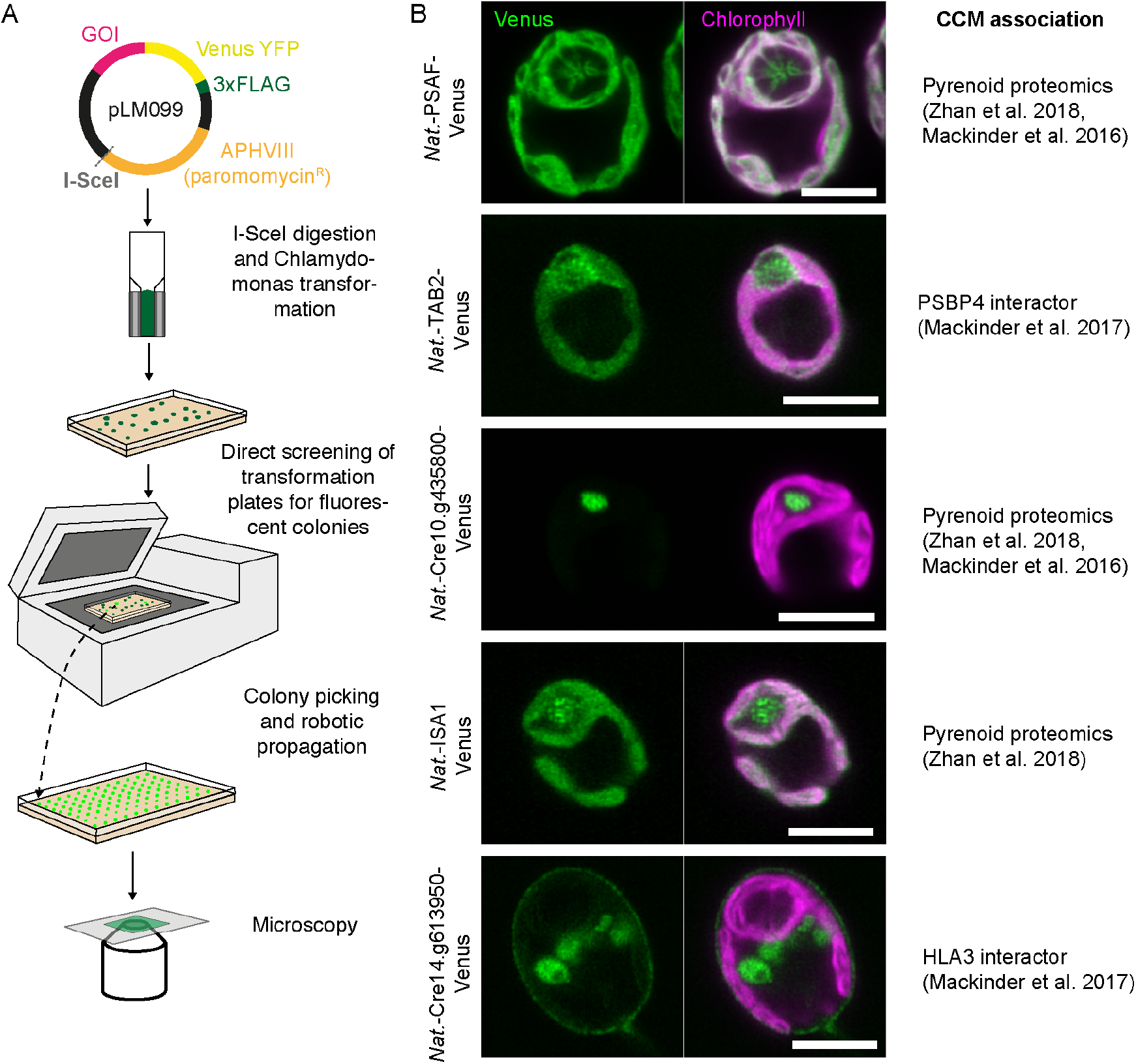
Transformation and localization of a subset of recombineered targets. **A** Chlamydomonas transformation pipeline. The I-SceI cut site allows vector linearization prior to Chlamydomonas transformation via electroporation. Transformants are directly screened for fluorescence using a Typhoon scanner (GE Healthcare) and then picked and propagated prior to imaging. **B** The localization for a subset of the recombineered target genes. Localizations agree with data from an affinitypurification followed by mass spectrometry study (Mackinder et al. 2017) or pyrenoid proteomics (Zhan et al. 2018 and/or Mackinder et al. 2016). Scale bars: 5 μm.

Cre10.g435800 localized to the pyrenoid matrix, and analysis of the translated product of Cre10.g435800 shows that it belongs to a family of NAD-dependent epimerase/dehydratases (IPR001509) and contains a UDP-galactose 4-epimerase domain that may be involved in galactose metabolism. Its role in pyrenoid function is unclear. Localization of ISA1 shows it was enriched in the pyrenoid with an uneven distribution. ISA1 is a starch debranching enzyme that is essential for starch synthesis with *ISA1* deletion lines lacking both chloroplast and pyrenoid starch (Mouille et al., 1996). The presence of pyrenoid starch and its correct organization is critical for correct CCM function (Itakura et al., 2019; Toyokawa et al., 2020), with the absence of starch in an *ISA1* knock out (4-D1) having incorrect LCIB localization (see below), retarded growth at very low CO_2_ (0.01% v/v) and reduced inorganic carbon affinity (Toyokawa et al., 2020). Interestingly in Toyokawa et al. (2020) they failed to attain localization data for an ISA1-mCherry fusion driven by the HSP70A/RBCS2 hybrid promoter.

Cre14.g613950 encodes a protein belonging to the ABC transporter family identified as an interactor of HLA3 (high light activated gene 3) (Mackinder et al., 2017), a putative HCO_3_^-^ transporter located in the plasma membrane (Duanmu et al., 2009; Gao et al., 2015). Like HLA3, Cre14.g613950 shows a typical plasma membrane localization pattern with YFP signal at the cell periphery and signal typical of the Golgi network.

### Development of backbones with additional tags and markers

To further expand the functional application of our recombineering pipeline we designed additional backbone vectors that enable protein tagging with the fluorophores mScarlet-i (Bindels et al., 2017), mNeonGreen (Shaner et al., 2013) and mTurquoise2 (Goedhart et al., 2012) and that allow selection with hygromycin or zeocin (Figure 5A). This enables complementation of Chlamydomonas Library Project (CLiP) mutants that have been generated using the *AphVIII* marker conferring paromomycin resistance (Li et al., 2019) and also enables expression of two or three differently tagged proteins within the same cell. For comparison, we tested these vectors on *LCI9* (Cre09.g394473), which encodes the low-CO_2_ inducible protein LCI9 that, via PCR-based cloning, we previously showed to localize to the pyrenoid periphery (Mackinder et al., 2017). Recombineered *LCI9* was 7160 bp long including the native promoter region. All fluorophores displayed the same pyrenoid periphery localization pattern (Figure 5B) and agree with the localization information obtained when LCI9 expression was driven from the *PSAD* promoter (Mackinder et al., 2017), thus further supporting that the use of ~2000 bp upstream regions as promoters for transgene expression is tenable.

**Figure 5.**
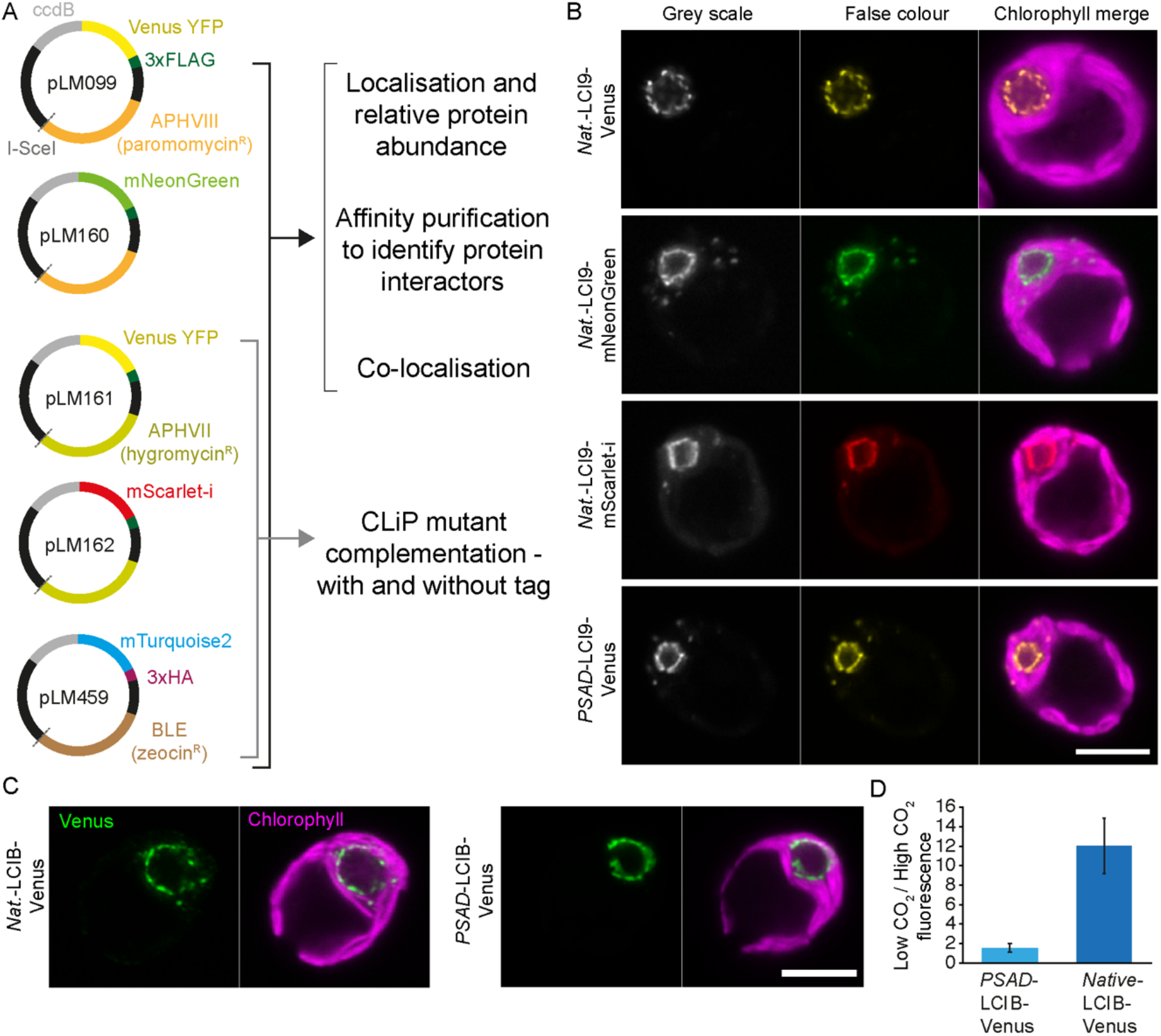
Development and application of different recombineering vectors to enable novel biological insights into Chlamydomonas biology. **A** Additional vectors for tagging with different fluorophores and for complementation of Chlamydomonas library mutants generated using insertion of the *AphVIII* paromomycin resistant gene. **B** Localization of LCI9 with different fluorescence protein tags. *LCI9* was recombineered with its native promoter (*nat*.) using pLM099, pLM160 and pLM162. A previously developed line cloned by PCR and using the constitutively expressed promotor *PSAD* is shown for comparison (*PSAD*-LCI9-Venus). Scale bar: 5 μm. **C** A comparison of the low CO_2_ upregulated gene LCIB cloned with its native promoter via recombineering vs LCIB under the constitutive *PSAD* promoter. Cells were grown and imaged at atmospheric CO_2_ levels. Scale bar: 5 μm. **D** Relative change in LCIB-Venus fluorescence between high (3% v/v) and low (0.04% v/v) CO_2_ when expressed from the constitutive *PSAD* promoter vs expression from the native LCIB promoter.

To further confirm that localization of proteins driven by their native promoter does not differ from those driven by the constitutive *PSAD* promoter we compared localization between native-LCIB-Venus and *PSAD*-LCIB-Venus. LCIB is an essential CCM component that shows dynamic relocalization to the pyrenoid periphery at CO_2_ levels <0.04% (Yamano et al., 2010). LCIB expressed from its endogenous promoter was localized to the pyrenoid periphery at very low CO_2_ (0.01% v/v), in full agreement with localization data when LCIB expression is driven by the constitutive *PSAD* promoter (Figure 5C).

### Maintaining the native promoter enables relative protein abundances to be monitored

As our pipeline retains the native promoter of the target gene we hypothesized that fluorescence output would be representative of relative protein abundance. To test this we grew lines with LCIB driven from either the constitutive *PSAD* promoter (*PSAD-*LCIB-Venus) or its native promoter (*Native*-LCIB-Venus). LCIB-Venus signal stayed constant between high (3% v/v) and low (0.04% v/v) CO_2_ when LCIB was expressed from the *PSAD* promoter (*PSAD-*LCIB-Venus), but showed an approximate 12-fold increase between these conditions when the native promoter was used (Figure 5D). This agrees with previous immunoblotting data, in which a >10-fold increase was seen in LCIB abundance when cells were transferred from high CO_2_ to low CO_2_ (Yamano et al., 2010). This indicates that our recombineering lines can be used to monitor relative protein abundance across different growth conditions.

## Discussion

We have established a rapid recombineering based method to clone large and complex Chlamydomonas genes from BACs. Our approach circumvents the challenges associated with cloning large, GC-rich and complex genes that are prevalent in Chlamydomonas. We demonstrate that the method can be applied for small batch cloning as well as 96-well high-throughput cloning. Our overall cloning success rate (combined batch and high-throughput results) was 77%, considerably higher than our previous PCR-based high-throughput cloning pipeline (48%), which was considerably inflated due to an enrichment of small target genes. Our overall success rate is slightly lower when compared to recombineering pipelines in other organisms, with success rates of 89% achieved in *C. elegans* (Sarov et al., 2012) and ~93% for Arabidopsis (Brumos et al., 2020). Our comparable success rate of 92% for our batch cloning approach indicates that our reduced overall efficiency could be further improved by additional optimization of our high-throughput pipeline. It is also likely that this reduced overall efficiency is due to the complexity of the Chlamydomonas genome (Figure 1), with DNA secondary structure linked to recombineering failure (Nelms and Labosky, 2011).

To enable expression of multiple fluorophores simultaneously and for the complementation of CLiP mutants we designed a series of vectors with modern fluorophores and varying selection markers and demonstrated their performance in Chlamydomonas (Figure 5). The presence of either 3xFLAG or 3xHA tag enables the vectors to be used for affinity purification to explore interacting partners of tagged proteins. Different fluorophore pairs (i.e. mNeonGreen and mScarlet-i) could also be used for FRET based studies to explore protein-protein interactions. In addition, all vectors can be used for cloning genes without fluorescence tags or with just short affinity tags (3xFLAG and 3xHA).

Due to the size independence of our method we could maintain the native promoter of target genes. For two genes, LCI9 and LCIB, the comparison between native promoter-driven expression and *PSAD* promoter-driven expression showed no noticeable differences in localization. Interestingly, using a native promoter allows relative protein abundance to be tracked between conditions (Figure 5D). Once validated, acquiring relative abundance data is straightforward and can be easily parallelized. This enables relative protein abundance to be tracked in real-time under a broad range of conditions. Future experiments could include tracking relative protein abundance in 96-well libraries of tagged proteins in response to a perturbation (i.e. high to low CO_2_ transition). This would be highly supportive of available transcriptomic and proteomic data sets and provide novel insights into cellular processes (Mettler et al., 2014; Zones et al., 2015; Strenkert et al., 2019). Although our relative abundance data for LCIB appears to closely reflect immunoblotting data, it should be noted that using a native promoter may not always fully reflect native changes. This discrepancy can be due to insertional effects caused by integration into transcriptionally unfavourable regions of the genome and absence of cis-regulatory regions in the recombineered construct,or transcriptional silencing (Schroda, 2019). At a protein level, fluorescent protein folding time could affect protein stability and turnover and the presence of the fused fluorescence protein could affect function or multi-subunit assembly.

Whilst our approach allows the native promoter and 5’UTR region to be cloned, the native 3’UTR is not maintained. This could be addressed through a two-step recombineering pipeline where the tag is first inserted into the BAC at the desired location, markers could then be removed via a *Flp-FRT* recombinase system (Sarov et al., 2006; Brumos et al., 2020), and the edited target gene can then be retrieved into a final Chlamydomonas expression vector. When establishing our pipeline, we decided not to pursue this strategy in order to maximise the success rate by limiting the number of steps, with a focus on developing a simple, easy to apply approach. In addition, whilst we have focused on C-terminal tagging as this allows conservation of N-terminal transit peptides required for organelle targeting, our recombineering pipeline could be applied for N-terminal tagging by modification of our cloning vectors with a constitutive promoter and N-terminal tag.

One limitation we encountered was that only 86% of nuclear genes are covered by the BAC library. However, this value only takes into account ~73% of BACs, with the remaining BACs potentially incorrectly mapped to the current version of the Chlamydomonas genome (see Supplemental Method 2B). Our analysis suggests the true percentage of genes covered could be higher than 86% but confirming this may require a careful re-mapping of the library. Another potential solution could be cloning from fosmids, with a Chlamydomonas fosmid library becoming available in the near future from the Chlamydomonas Resource Centre. The use of fosmids, with smaller DNA fragments compared to BACs, could help improve efficiency by reducing off-target recombination between the PCR-amplified cassette and the BAC or by reducing recombination between two repetitive regions of the BAC.

Our recombineering approach has enabled the efficient cloning of large and complex genes that could not be achieved via PCR-based cloning. It opens the door to a better understanding of the functional role of a large fraction of the Chlamydomonas genome though protein localization, protein-protein interaction studies, real-time monitoring of relative protein abundance and complementation of mutants. In addition, it provides a highly complementary method to the recently released CLiP mutant collection.

## Methods

### Plasmid and cassette construction

Fragments for pLM099 were amplified by PCR (Phusion Hotstart II polymerase, ThermoFisher Scientific) from the following plasmids: pLM005 (Mackinder et al., 2017) for Venus-3xFLAG, PSAD terminator and *AphVIII*; pNPC2 for the p15A origin of replication; pLM007 for the *Kan^R^* resistance gene; Gateway pDONR221 Vector (ThermoFisher Scientific) for the counter-selection *ccdB* gene. The resulting amplicons were gel purified (MinElute Gel Extraction Kit, QIAGEN) and assembled by Gibson assembly (see Figure S1 for detailed map). pLM160 was constructed from pLM099 to replace CrVenus with mNeonGreen (Shaner et al., 2013), and pLM161 was constructed from pLM099 to replace the paromomycin resistance gene (*AphVIII*) with the hygromycin resistance gene (*AphVII*). pLM162 was constructed from pLM161 with the synthetic fluorophore mScarlet-i (Bindels et al., 2017) replacing CrVenus. pLM459 was constructed from pLM161 to replace CrVenus with mTurquoise2 (Goedhart et al., 2012), the 3xFLAG with the 3xHA haemagglutinin tag, and *AphVII* with the zeocin resistance gene (*BLE*). Gene-specific cloning primers were designed to amplify a ~4.6 kbp cassette from the recombineering vectors pLM099, 160, 161, 162 and 459 (Figure 5), excluding *ccdB*, and providing 50 bp of sequence homology to the target gene an average of ~2500 bp upstream of the 5’UTR and directly upstream of the stop codon. This enables the retrieval of each target gene into the cassette in frame with a fluorescent tag and with the native promoter region intact. All oligonucleotide and plasmid sequences can be found in Supplemental Data Sets 3 and 4.

### Culturing

*E. coli* cells were cultured in lysogeny broth (LB) or yeast extract nutrient broth (YENB) at 37°C unless they contained the temperature sensitive pSC101-BAD-gbaA-tet (pRed), in which case 30°C was used. All DNA for transformation was introduced by electroporation and transformants were recovered in super optimal broth with catabolite repression (SOC). DH10B cells containing fragments of the Chlamydomonas genome in the form of BACs were obtained from the Clemson University Genomics Institute (now distributed by the Chlamydomonas Resource Centre, University of Minnesota, USA). DB3.1 cells expressing the *ccdB* antidote gene, *ccdA*, were obtained from ThermoFisher Scientific and used for maintenance of the recombineering vectors.

Chlamydomonas wild type cells (strain CC-4533) were cultured in Tris-acetate-phosphate media (TAP) with revised Hutner’s trace elements (Kropat et al., 2011). Assembled recombineering vectors were prepared for transformation into Chlamydomonas by restriction digest with I-SceI endonuclease (NEB). Transformation and selection were performed in accordance with Mackinder et al. (2017) using a Typhoon Trio fluorescence scanner (GE Healthcare). Viable Chlamydomonas transformants were screened for CrVenus and mNeonGreen expression at 555/20 nm, and for mScarlet-i at 615/12 nm. Several strains emitting the strongest fluorescence for each line were picked and cultured in Tris-phosphate minimal media, then imaged by fluorescent microscopy to visualise protein localization.

### Microscopy

Sample preparation for microscopy was performed as per (Mackinder et al., 2017). Images were acquired using a Zeiss LSM880 confocal microscope on an Axio Observer Z1 invert, equipped with a 63x 1.40 NA oil planapochromat lens. Images were analysed using ZEN 2.1 software (Zeiss) and FIJI. Excitation and emission filter settings were as follows: Venus and mNeonGreen, 514 nm excitation, 525-550 nm emission; mScarlet-i, 561 nm excitation, 580-600 nm emission; and chlorophyll, 561 nm excitation, 665-705 nm emission.

### Recombineering procedure for 1-step subcloning and tagging

The following outlines the batch-scale recombineering protocol. Extended batch and multi-well plate-scale recombineering protocols are supplied in Supplemental Method 1.

For each target, a recombineering cassette was amplified from plasmid pLM099 (Phusion Hotstart II polymerase, ThermoFisher Scientific) using primers containing 50 bp homology arms, one homologous to a region upstream of the annotated start codon of the target gene, and one homologous to the 3’ end of the coding sequence (excluding the stop codon). The resulting PCR product was purified (MinElute Gel Extraction Kit, QIAGEN) and its concentration measured using a nanodrop spectrophotometer. Upstream region lengths ranged from 1000-4000 bp from the start codon, with an average of ~2500 bp. For two genes, Cre04.g220200 and Cre16.g678661, the first 50 bp of the 5’UTR was used as the upstream homology region due to BAC coverage limitations.

The pRed plasmid, pSC101-BAD-gbaA-tet, was extracted from *E. coli* cells grown overnight at 30°C (Plasmid Mini Kit, QIAGEN), and its concentration measured by nanodrop. *E. coli* cells harbouring a BAC containing the target gene were recovered from the Chlamydomonas BAC library and used to inoculate 20 ml of YENB media containing 12.5 μg/ml chloramphenicol, followed by overnight growth in a 50 ml conical flask at 37°C with vigorous shaking. After 16 h of growth, 120 μl of the culture was used to inoculate 4 ml of fresh YENB containing 12.5 μg/ml chloramphenicol. This was grown for ~2 h at 37°C until an optical density (OD_600_) of 2 was reached. 2 ml of the culture was then incubated on ice for 2 min, followed by centrifugation at 5000 x g for 10 min at 4°C. After removing the supernatant, the pellet was placed back on ice and washed by resuspension in 1 ml of chilled 10% glycerol, followed immediately by centrifugation at 5000 x g for 10 min at 4°C. The resulting supernatant was removed, and the pellet was placed back on ice and resuspended in 100 μl of 0.1 ng/μl pRed. This mixture was transferred to a pre-chilled 2 mm gap electroporation cuvette and electroporated at 2500 V, 400 Ω and 25 μF using a Gene Pulser II (Bio-Rad). The electroporated cells were immediately recovered in 800 μl SOC and incubated at 30°C for 90 min with vigorous shaking. The whole outgrowth was added to 20 ml of YENB containing 12.5 μg/ml chloramphenicol and 5 μg/ml tetracycline and grown overnight at 30°C with vigorous shaking.

After 16 h of growth, 600 μl of culture was used to inoculate 4 ml of fresh YENB containing 12.5 μg/ml chloramphenicol and 5 μg/ml tetracycline. This was grown for 3 h at 30°C, or until reaching an OD_600_ >2, at which point 80 μl of 10% L-arabinose was added to induce pRed expression and growth was shifted to 37°C for 1 h with vigorous shaking. 2 ml of the induced culture was incubated on ice for 2 min, then centrifuged at 5000 x g for 10 min at 4°C, the supernatant removed, and the pellet placed back on ice. Cells were then washed in 10% glycerol, centrifuged at 5000 x g for 10 min at 4°C, the supernatant removed, and the pellet placed back on ice. The pellet was resuspended in 100 μl of 5 ng/μl PCR product and transferred to a pre-chilled 2 mm gap electroporation cuvette, followed by electroporation as before. Electroporated cells were immediately added to 800 μl of SOC and recovered at 37°C for 90 min with vigorous shaking. 450 μl of outgrowth was spread onto 1.5% LB-agar containing 25 μg/ml kanamycin, airdried and incubated overnight at 37°C. Selected colonies were used to inoculate 4 ml of LB containing 25 μg/ml kanamycin and grown for 16-18 h at 37°C with shaking. Recombineering products were extracted and validated by restriction digest using appropriate enzymes, followed by Sanger sequencing using primers designed to amplify the junctions between the pLM099-derived cassette and the target region.

### Genome analysis

Chlamydomonas, Arabidopsis, yeast and wheat nuclear genes were analysed for gene length and sequence complexity. In this work, the complexity of a sequence is defined in relation to intron prevalence, GC content, and the prevalence of various repeat regions. We designate regions containing a high frequency of repeats as being more complex than regions with a low frequency. This reflects the increased potential for cloning complications presented by sequences with large numbers of repetitive regions, though it differs from the descriptions given by Morgulis et al (2006). Sequences were analysed for complexity using the freely available bioinformatics software detailed below (see Supplemental Method 3 for settings), and outputs were processed using custom python scripts (Supplemental Code; see Supplemental Method 4 for usage information). GC content was calculated such that unannotated bases had no effect on the value.

#### Sequence data sources

Unspliced nuclear gene sequences for *Chlamydomonas reinhardtii* (version 5.5) were obtained from Phytozome Biomart. Sequence data for *Arabidopsis thaliana* (TAIR10 assembly) and *Triticum aestivum* nuclear genes (International Wheat Genome Consortium assembly) were obtained from EnsemblPlants BioMart. Analysis was limited to the 105,200 chromosome-assigned wheat genes. Sequence data for *Saccharomyces cerevisiae* (S288C reference genome, 2015 release) were obtained from the Saccharomyces Genome Database. Gene sequences were appended to include all annotated UTRs and introns, resulting in a dataset that is more closely comparable to the unspliced gene data used for Chlamydomonas, Arabidopsis and wheat.

#### Analysis of repeats

Repetitive regions in the nucleotide sequences analysed in this work are categorized into simple repeats and global repeats. We use the term simple repeats to refer to relatively short (tens to hundreds of bases) repetitive regions in a nucleotide sequence that display regular or semi-regular repeating patterns. We include consecutive repeating motifs of varying unit lengths, known as tandem repeats, as well as inverted patterns in which a short region is followed closely (or immediately, if palindromic) by its reverse complement sequence. Chlamydomonas genes were analysed for tandem repeats using Tandem Repeats Finder (Benson, 1999). The default settings were modified to provide a cutoff for detection such that no repeats under 10 bp in length were reported (see Supplemental Method 3A). All Tandem Repeats Finder outputs were processed using a custom python script and analysed in spreadsheet format to generate mean values for the number of genes with either, (1) at least one mononucleotide repeat ≥10 bp in length and with ≥90% identity; (2) at least one di- or trinucleotide repeat ≥20 bp in length with ≥90% identity; (3) at least one tandem repeat ≥20 bp in length, with a period length of four or more (tetra+), with ≥90% identity; and (4) the mean number of repeats of these types per kilobase of sequence.

Chlamydomonas genes were analysed for inverted repeats using the Palindrome Analyser webtool (Brázda et al., 2016), available at http://bioinformatics.ibp.cz:9999/#/en/palindrome. The default settings were modified to report repeats with a maximum of 1 mismatch for every 10 bp of stem sequence, a maximum spacer length of 10 bp and a maximum total length of 210 bp (see Supplemental Method 3B for settings). All Palindrome Analyser outputs were downloaded and analysed in spreadsheet format to generate mean values for the number of genes containing one or more inverted repeats over 20 bp long with ≥90% identity, and the mean number of inverted repeats of this type per kilobase.

All nuclear genes from Chlamydomonas (Figure 1B), Arabidopsis, yeast and wheat (Figure 1F), and recombineering target regions (Figure 3B and C) were analysed for global repeats using the NCBI WindowMasker program (Morgulis et al., 2006). We use the term global repeats to denote the combined number of individual masked regions detected by the WindowMasker modules DUST and WinMask. DUST detects and masks shorter repetitive regions including tandem and inverted repeats, overlapping with and providing support for the Tandem Repeats Finder and Palindrome Analyser outputs. WinMask detects and masks families of longer repetitive regions that do not necessarily occur adjacently in the genome. Default settings were used throughout (see Supplemental Method 3C). These modules mask repetitive regions using only the supplied sequence as a template.

#### uORFs, transcripts and intron analysis

Data on the presence of uORFs in Chlamydomonas transcripts were obtained from the results of a BLASTP analysis performed by Cross (2015) and adapted to provide the per-gene values. A list of Chlamydomonas transcripts was downloaded from Phytozome Biomart and used to identify the number of genes with more than one transcript model. Genomic data detailing the number and order of exons within each gene were also downloaded from Phytozome Biomart; this information was used to ascertain the number of genes containing introns in their translated and untranslated regions.

#### Primer analysis

To assess the impact of inefficient priming on PCR-based cloning, analysis was performed on a dataset of PCR primers designed to clone every gene in the Chlamydomonas genome from start to stop codon using gDNA as the template, and generated such that the predicted Tm difference for each pair was not more than 5°C where possible. Primer sequences were then assessed against four thresholds pertaining to efficient priming, set in accordance with advice found in the Primer3 manual, support pages provided by IDT, and the Premier Biosoft technical notes. These thresholds relate to primer length,propensity for secondary structure formation, the presence of repeats and the GC content of the 3’ end. Long primers can have a reduced amplification efficiency, secondary structure formation can reduce the number of primers available to bind to the intended template during a PCR, multiple repeats can increase the risk of mispriming, and a high 3’ end GC content can increase the risk of primer-dimer formation. Thresholds for each were set as follows: (1) primer length should not be more than 30 bp, (2) the Δ*G* required to disrupt predicted secondary structures should be above −9 kcal/mol at 66 or 72°C, (3) tandem single nucleotides or dinucleotide motifs should repeat no more than 4 times, and (4) the 3’ end should consist of no more than 4 G/C bases in a row. The number of primers in breach of each of these thresholds is shown in Figure 1D as a percentage of the dataset. The percentage of unsuitable primer pairs was calculated by counting pairs for which one or both primers breached one or more of these thresholds. Tm considerations were omitted from analysis since Chlamydomonas genes have an unusually high GC content, so primers designed to amplify gDNA are expected to have higher than recommended Tms according to generic primer design guidelines. GC content was calculated such that unannotated bases had no effect on the value.

To complement these results, primers were analysed using the check_primers algorithm from Primer3 (Rozen and Skaletsky, 2000). Settings used were as default for Primer3Plus (Untergasser et al., 2007) – an updated, online version of the Primer3 package – with minimal modifications that included removing the Tm constraints (see Supplemental Method 3D for full settings used). The output was analysed with a custom python script that reported the primary reason for rejection of individual primers (see Supplemental Method 4C). Tm was removed as a constraint to allow for more detailed analysis of primer sequence parameters, since the default maximum allowable Tm for Primer3Plus is 63°C, which results in rejection of almost 90% of primers for this reason alone if used. 1.6% of primers were too long to be considered for analysis (>36 bp); these were included in Figure 1D (orange bar) as having been rejected for breaching the length constraint. The majority of rejected primers produced one of the following three reasons for rejection: (1) ‘high end complementarity’ for primer pairs, which implies a high likelihood that the 3’ ends of the forward and reverse primers will anneal, enabling amplification of a short, heterogeneous primer-dimer (cross-dimer); (2) ‘high end complementarity’ for single primers, which implies a high likelihood that a primer’s 3’ end will bind to that of another identical copy, self-priming to form a homogenous primer-dimer (self-dimer); and (3) ‘high any complementarity’ for single primers, which implies a high likelihood of self-annealing without necessarily self-priming, relevant to both the inter-molecular annealing of identical copies and to instances of hairpin formation resulting from intra-molecular annealing. Primers rejected for these three reasons are labelled in Figure 1D (orange bar) as cross-dimers, self-dimers and hairpins, respectively.

### BACSearcher python resource

Suitable BACs containing the target genes were identified using a python script that also identifies 50 bp binding sites for recombineering cloning primers and provides sequences for primers that can be used to check for the presence of a target gene within a BAC (see Supplemental Method 2). BACSearcher output is available for all 17,741 genes in the genome in Supplemental Data Set 1. For individual targets in our recombineering pipeline that were not covered by a BAC in the BACSearcher output, an alternative method was employed to search for BAC coverage. This method is detailed in Supplemental Method 2, along with usage and modification instructions for BACSearcher.

### Data and software availability

The python computer code used for identifying BACs and suitable homology regions for recombineering is available at https://github.com/TZEmrichMills/Chlamydomonas_recombineering and supplied as Supplemental Code.

## Author Contributions

TEM developed the initial recombineering pipeline, designed and assembled the original pLM099 recombineering plasmid and performed the genome wide analysis. GY, TEM and TKK assembled additional recombineering plasmids. TEM and GY optimized and performed the large-scale recombineering pipeline. GY performed the microscopy and Venus quantification data. JB, IG, CSL, CEW and TKK supported the development and implementation of the recombineering pipeline. JWD wrote the BACSearcher code and provided bioinformatics support to TEM. LCMM conceived the idea and led the research. LCMM and MPJ received funding to support the work. LCMM, TEM and GY wrote the manuscript.

## Supplemental Data

**Figure S1.**
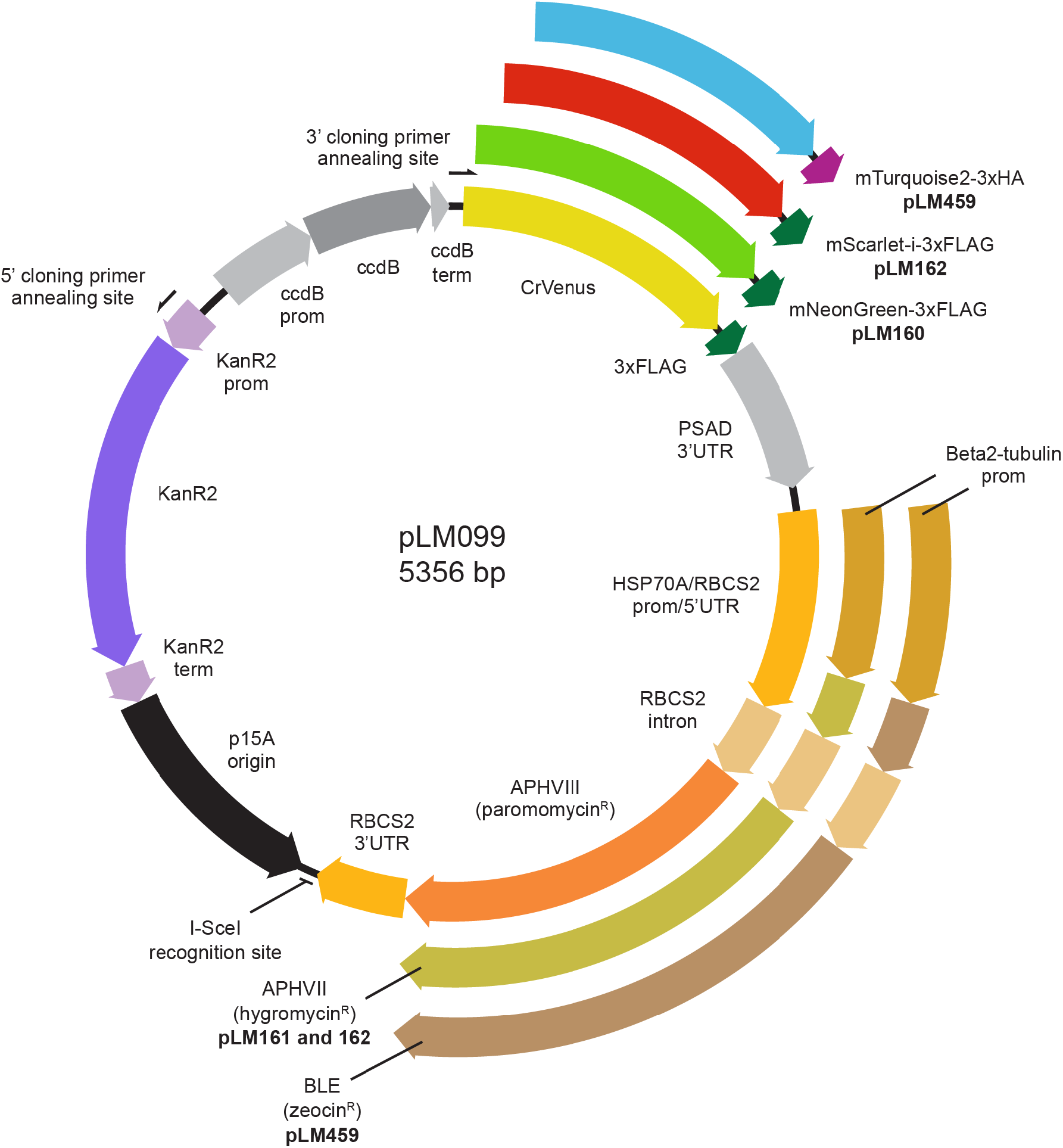
Plasmid map for pLM099 and derivative recombineering vectors. PCR amplification with 5’ and 3’ cloning primers at the annealing sites shown results in a ~4.6 kbp linear cassette for recombineering target genes in-frame with a fluorescent protein and affinity tag. For each recombineering vector, the fluorescent protein sequence is preceded by a short, flexible linker (GGLGGSGGR) and followed by a tri-glycine linker prior to the affinity tag. The PSAD 3’UTR terminates all four fluorescent protein-affinity tag cassettes. The RBCS2 3’UTR terminates all three Chlamydomonas selection cassettes. The same RBCS2 intron is present in all three Chlamydomonas selection cassettes, but is only intra-exonic in the hygromycin and zeocin resistance cassettes.

**Figure S2.**
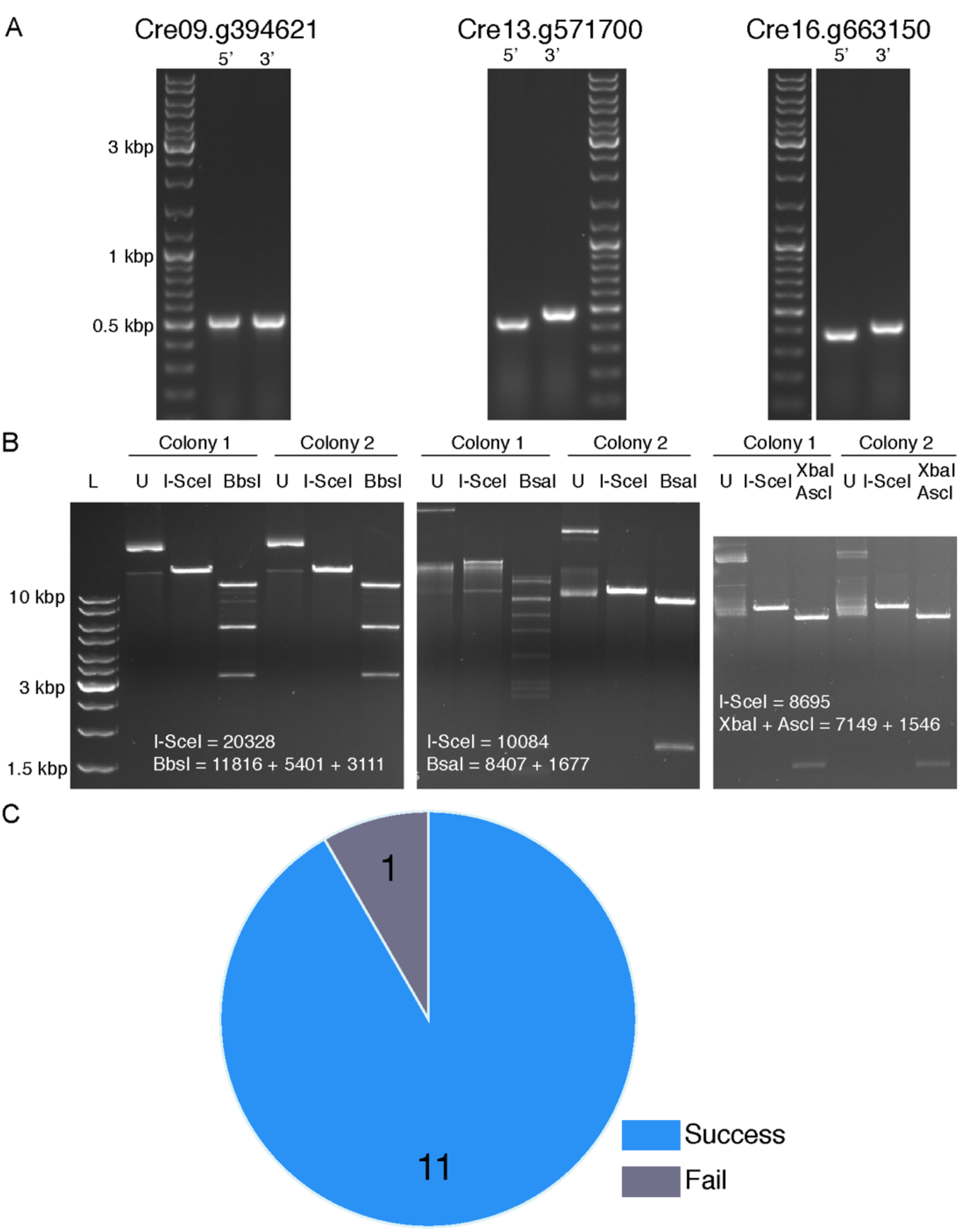
Batch-scale recombineering results. **A** Three examples of colony PCRs to check for presence of target genes in BACs. Primer pairs were designed to the 5’ and 3’ end of each target gene. All amplicons were of the expected size. **B** Restriction digest checks for isolated recombineered plasmids from two colonies per gene, corresponding to the same genes as in **A**. Expected sizes are shown in bp. Note that colony 1 for Cre13.g571700 gives the incorrect size and banding patterns after digestion indicating incorrect recombination. U: undigested. **C** Overall batch-scale recombineering success for 12 target genes.

Supplemental Method 1. Protocols for batch- and large-scale recombineering

Supplemental Method 2. BACSearcher usage

Supplemental Method 3. Bioinformatics software usage

Supplemental Method 4. Bioinformatics python analysis

Supplemental Data Set 1. BACSearcher output

Supplemental Data Set 2. Large-scale pipeline results summary

Supplemental Data Set 3. Oligonucleotide sequences

Supplemental Data Set 4. Plasmid sequences

Supplemental Code. BACSearcher python code, BACSearcher precursor files and python code for processing outputs from bioinformatics programs

## Acknowledgments

This work was funded by the UK Biotechnology and Biological Sciences Research Council Grant BB/R001014/1 (to LCMM); Leverhulme Trust Grant RPG-2017-402 (to LCMM); BBSRC DTP2 BB/M011151/1 (to TEM and MPJ); BBSRC DTP2 BB/M011151/1a (to JB and LCMM); University of York Biology Pump Priming award (to LCMM); and University of York Biology Start-up grant (to LCMM). We would like to thank Guy Mayneord for programming advice, the University of York Biosciences Technology Facility for confocal microscopy and bioinformatics support, and Mihail Sarov (University of Dresden, Germany) for technical advice and providing plasmids pSC101-BAD-gbaA-tet and pNPC2.

## Notes

### Competing Interest Statement

The authors have declared no competing interest.

