## Supplementary material for "A recombineering pipeline to clone large and complex genes in Chlamydomonas": Supplemntal Code: BACSearcher.txt

#!/usr/bin/env python3

"""
BACSearcher
@author John W. Davey
Associated GitHub repository: https://github.com/TZEmrichMills/Chlamydomonas_recombineering
For additional usage information, see Supplemental Method 2 or README.md in the repository.

Usage:

BACSearcher can be initiated from the command line using options -p, -f, -g, -l, -w and -o.
Modifications can be made to the default parameters using additional options -q, -r, -s, -t, -u, and -v.
Default usage example:

./BACSearcher.py -p BACs_fosmids.pairs.tsv -f Creinhardtii_281_v5.0.fa.gz -g Creinhardtii_281_v5.5.gene.gff3.gz -l Gene_shortlist.txt -w Bac_wells.txt -o Chlamydomonas_BACSearcher_results

BACs_fosmids.pairs.tsv refers to precursor file I (below); 
Creinhardtii_281_v5.0.fa.gz refers to precursor file II; 
Creinhardtii_281_v5.5.gene.gff3.gz refers to precursor file III; 
Gene_shortlist.txt refers to precursor file IV; and
BAC_wells.txt refers to precursor file V.
Chlamydomonas_BACSearcher_results is an output file name. 

Precursor files:

Precursor file I: TSV file containing the coordinates of the start and end of each valid BAC in the library. BACs are included as valid if their start and end sequences are mapped to the same chromosome and are in the correct orientation, (i.e., one end on each strand).
Precursor file II: Zipped FASTA file (.fa.gz extension) containing the gene sequences for all Chlamydomonas nuclear genes.
Precursor file III: Zipped GFF file (.gff3.gz extension) containing version 5.5 annotation information for the Chlamydomonas genome.
Precursor file IV: (Optional) TXT file containing the Cre IDs for all genes of interest to be processed, one per line, each appended with ‘.v5.5’. If this file is not provided, BACSearcher will process all nuclear genes and produce a TSV file of the results with the name specified by -o (see Example usage, above).
Precursor file V: TXT file containing the plate and well coordinates of each BAC in the library, in the format ‘A-B-C’, where A is the plate number, B the row number and C the column number. This file is provided in the Supplemental Code ZIP folder as BAC_wells.txt and is also available from the GitHub repository (see Code availability section).
Precursor file VI: (Optional) DB file generated from III during the first running of the script, which can be used in place of III in future runs. 

I and V are provided as BACs_fosmids.pairs.tsv and BAC_wells.txt in the Supplemental Code ZIP folder, as well as in the associated GitHub repository. 
II and III are available for download from Phytozome V12 entitled Creinhardtii_281_v5.0.fa.gz and Creinhardtii_281_v5.5.gene.gff3.gz. 

Note: When supplied with a GFF file via -g, BACSearcher will generate a gffutils database for the GFF file (precursor file VI). 
BACSearcher can also use this database directly with the -d option, saving the effort of regenerating the database.

Modifications to the defaults:

-q followed by an integer, x, will change the default lengths of the reported 5’ homology regions to an integer, x bp.
-r followed by an integer, x, will change the default length of the 3’ homology region to an integer, x bp.
-s followed by an integer, x, will direct the script to search for suitable homology regions in the 1000 bp downstream of an upstream position, x bp. For example, if x=5000 the script will search for regions between 4000 and 5000 bp upstream of the start codon of each gene.
-u will direct the script to search for homology regions in the region defined by -s but measured from the 5’UTR instead of the start codon. 
-t followed by an integer, x, will direct the script to report only those BACs that cover x bp downstream from the stop codon of each gene.
-v will direct the script to report BACs that cover a downstream region defined by -t but measured from the 3’UTR instead of from the stop codon of each gene. 


"""

import argparse, os, sys, gzip
import gffutils
from collections import defaultdict
from intervaltree import Interval, IntervalTree
from Bio import Seq
from Bio import SeqIO
from Bio import SeqUtils
from Bio.SeqUtils import GC
from Bio.Seq import Seq
from primer3 import designPrimers

parser=argparse.ArgumentParser(description='''
    -g genegff
    -d genedb
    -p plasmidpairs
    -l genelist
    -o outputstub
    -f genome FASTA
    -w wells
    -n num_results
    -q fiveprime_homology_length
    -r threeprime_homology_length
    -s fiveprime_flank length
    -t threeprime_flank_length
    -u fiveprime_utr_flank
    -v threeprime_utr_flank
    ''')

parser.add_argument('-p', '--plasmidpairs', type=str, required=True)
parser.add_argument('-f', '--genomefasta', type=str, required=True)
parser.add_argument('-w', '--wells', type=str, required=True)
parser.add_argument('-g', '--genegff', type=str)
parser.add_argument('-d', '--genedatabase', type=str)
parser.add_argument('-l', '--genelist', type=str)
parser.add_argument('-n', '--num_results', type=int, default=5)
parser.add_argument('-q', '--fiveprime_homology_length', type=int, default=50)
parser.add_argument('-r', '--threeprime_homology_length', type=int, default=50)
parser.add_argument('-s', '--fiveprime_flank_length', type=int, default=3000)
parser.add_argument('-t', '--threeprime_flank_length', type=int, default=0)
parser.add_argument('-u', '--fiveprime_utr_flank', action='store_true')
parser.add_argument('-v', '--threeprime_utr_flank', action='store_true')
parser.add_argument('-o', '--outputstub', type=str, default="test")

args = parser.parse_args()

def load_annotation(gfffile, database):
    db_filename = None
    if database is not None:
        if os.path.isfile(database):
            db_filename = database
        else:
            print(f"Database {database} does not exist", file=sys.stderr)
    elif gfffile is not None:
        print(f"Loading annotation {gfffile} into database", file=sys.stderr)
        new_db_filename = gfffile + '.db'
        if os.path.isfile(new_db_filename):
            print(f"Annotation database {new_db_filename} already exists", file=sys.stderr)
        else:
            db = gffutils.create_db(gfffile, new_db_filename)
        db_filename = new_db_filename
    
    if db_filename is None:
        print("Please supply one of -g and -d for gene annotation and one of -a and -b for BAC annotation", file=sys.stderr)
        sys.exit()
    
    print(f"Loading annotation database {db_filename}")
    db = gffutils.FeatureDB(db_filename)
    
    return db, db_filename

def get_plasmid_locations(plasmidpairfile, length_threshold=300000):
    bacs = defaultdict(IntervalTree)
    fosmids = defaultdict(IntervalTree)
    with open(plasmidpairfile, 'r') as ppf:
        for line in ppf:
            if line.startswith("Plasmid"):
                continue
            plasmid, end1, end2, chromosome, start, end, length = line.rstrip().split('\t')
            start, end, length = int(start), int(end), int(length)
            if length > length_threshold:
                continue
            if plasmid.startswith('PTQ'):
                bacs[chromosome][start:end] = (plasmid, end1, end2, chromosome, start, end, length)
            elif plasmid.startswith('VTP'):
                fosmids[chromosome][start:end] = (plasmid, end1, end2, chromosome, start, end, length)

    return bacs, fosmids

class Plasmid:
    def __init__(self, plasmid_interval=None):
        if plasmid_interval:
            self.begin = plasmid_interval.begin
            self.end = plasmid_interval.end
            self.name = plasmid_interval.data[0]

            self.type = ''
            if self.name.startswith('PTQ'):
                self.type = 'BAC'
            elif self.name.startswith('VTP'):
                self.type = "Fosmid"

            self.ends = f"{plasmid_interval.data[1]},{plasmid_interval.data[2]}"
            self.length = plasmid_interval.data[6]
        else:
            self.begin = self.end = self.type = self.ends = self.name = self.length = ''
    
    def __repr__(self):
        return f"{self.name}\t{self.ends}\t{self.begin}\t{self.end}\t{self.length}"

class PrimerPair:
    def __init__(self, i, primer_results):
        self.i = i
        self.left_sequence  = primer_results[f'PRIMER_LEFT_{i}_SEQUENCE']
        self.right_sequence = primer_results[f'PRIMER_RIGHT_{i}_SEQUENCE']

        self.product_size   = primer_results[f'PRIMER_PAIR_{i}_PRODUCT_SIZE']
    
    def __repr__(self):
        return f"{self.left_sequence}\t{self.right_sequence}\t{self.product_size}"


def get_primers(flank, num_results=1):
    results=designPrimers(
        seq_args={
            'SEQUENCE_TEMPLATE': str(flank)
        },
        global_args={
            'PRIMER_NUM_RETURN':num_results,
            'PRIMER_PRODUCT_SIZE_RANGE':[150,250]
        }
    )

    primers = [PrimerPair(i, results) for i in range(results['PRIMER_PAIR_NUM_RETURNED'])]
    if len(primers) < num_results:
        for i in range(len(primers)-1, num_results):
            primers.append('\t\t')
    return primers

def mask_dinucleotides(flank):
    
    num_dinucs = 5
    flanklist = list(flank)
    for i in range(len(flank)-num_dinucs*2+1):
        dinuc = flank[i:i+2]
        if 'N' in dinuc:
            continue
        pos = [i, i+1]
        for j in range(i+2, len(flank), 2):
            if dinuc == flank[j:j+2]:
                pos.append(j)
                pos.append(j+1)
            else:
                break
        if len(pos) >= num_dinucs*2:
            for p in pos:
                flanklist[p] = 'N'
    flank = ''.join(flanklist)
    return flank

class Gene:
    def __init__(self, gene_name, gene_db, bacs, fosmids, wells, num_results, \
                 fiveprime_flank_length, threeprime_flank_length, \
                 fiveprime_utr_flank, threeprime_utr_flank, \
                 fiveprime_homology_length, threeprime_homology_length):

        self.gene_name = gene_name
        self.gene     = gene_db[self.gene_name]
        self.chromosome = self.gene.seqid
        self.strand = self.gene.strand
        self.num_results = num_results
        self.fiveprime_flank_length = fiveprime_flank_length
        self.threeprime_flank_length = threeprime_flank_length
        self.fiveprime_utr_flank = fiveprime_utr_flank
        self.threeprime_utr_flank = threeprime_utr_flank
        self.fiveprime_homology_length = fiveprime_homology_length
        self.threeprime_homology_length = threeprime_homology_length

        self.coding_start, self.coding_end, self.flank_start, self.flank_end = self.__get_location(gene_db)

        self.bacs = self.__get_plasmids(bacs, 'BAC')
        self.fosmids = self.__get_plasmids(fosmids, 'Fosmid')

        self.bac_wells = [wells[x.name[3:]] if x.name else '' for x in self.bacs]

        self.fiveprime_regions = self.__get_fiveprime_regions()

        self.threeprime_region = self.__get_threeprime_region()
        self.threeprime_gc = GC(self.threeprime_region)

        self.fiveprimers, self.threeprimers = self.__get_primers()

    def __get_location(self, gene_db):
        coding_start = coding_end = flank_start = flank_end = None

        for cds in gene_db.children(self.gene_name, featuretype="CDS"):
            if coding_start is None or cds.start < coding_start:
                coding_start = cds.start
            if coding_end is None or cds.end > coding_end:
                coding_end = cds.end
        
        start_anchor, end_anchor = coding_start, coding_end
        if self.fiveprime_utr_flank:
            start_anchor = self.gene.start
        if self.threeprime_utr_flank:
            end_anchor = self.gene.end

        if self.strand == '+':
            flank_start, flank_end = start_anchor - self.fiveprime_flank_length, end_anchor + self.threeprime_flank_length
        elif self.strand == '-':
            flank_start, flank_end = start_anchor - self.threeprime_flank_length, end_anchor + self.fiveprime_flank_length
        
        if flank_start < 1:
            flank_start = 1

        if flank_end > len(genome[self.chromosome]):
            flank_end = len(genome[self.chromosome])

        return coding_start, coding_end, flank_start, flank_end

    def __get_plasmids(self, plasmid_pairs, plasmid_type):
        # Sort intervals overlapping plasmid by length, shortest first
        # Return the first five plasmids that envelope the flanking region (defined in lines 183 and 185)
        plasmids = []
        plasmids_added = {}
        for plasmid_pair in sorted(plasmid_pairs[self.chromosome].overlap(self.flank_start, self.flank_end), key=lambda x:(x.length())):
            if plasmid_pair.begin < self.flank_start and plasmid_pair.end > self.flank_end and plasmid_pair.data[0] not in plasmids_added:
                plasmids.append(Plasmid(plasmid_pair))
                plasmids_added[plasmid_pair.data[0]] = 1

        plasmids = plasmids[:self.num_results]

        if len(plasmids) < self.num_results:
            for i in range(len(plasmids)-1, self.num_results):
                plasmids.append(Plasmid())

        return plasmids

    def __get_fiveprime_regions(self):
        region = upstream_pos = None
        
        flank = None
        # Extract 1000bp at the end of the flanking region (2-3kb away from gene start).
        if self.strand=='+':
            # If flank_start is 1, we want GFF positions 1-1000, Python coordinates 0-999, Python range 0:1000, which is flank_start-1:flank-start+999
            flank = genome[self.chromosome][self.flank_start-1:self.flank_start+999].seq
        elif self.strand=='-':
            # If flank_end is 1000, we want GFF positions 1-1000, Python coordinates 0-999, Python range 0:1000, which is flank_end-1000:flank_end
            flank = genome[self.chromosome][self.flank_end-1000:self.flank_end].reverse_complement().seq

        flank = mask_dinucleotides(flank)
        # Find the 50bp region in the 1000bp flanking region with minimum GC, then reverse complement for ease of addition to the 5' cloning primer
        low_gc_regions = sorted(
          [(i+2001, Seq(str(flank[i:i+self.fiveprime_homology_length])).reverse_complement()) 
             for i in range(len(flank)-self.fiveprime_homology_length)
             if 'N' not in flank[i:i+self.fiveprime_homology_length]], 
          key=lambda x:(GC(x[1]),x[0]))

        nonoverlaps = []
        for r in low_gc_regions:
            append = True
            for n in nonoverlaps: 
                if n[0]-(self.fiveprime_homology_length-1) < r[0] < n[0]+(self.fiveprime_homology_length-1):
                    append = False
            if append:
                nonoverlaps.append(r)

        if len(nonoverlaps) < self.num_results:
            for i in range(len(nonoverlaps), self.num_results):
                nonoverlaps.append(('',''))

        return nonoverlaps

    def __get_threeprime_region(self):
        if self.strand=='+':
            # Sequence is upstream of the end of the gene. Skip the stop codon (-3). We want the 50bp prior to the stop codon.
            # GFF is 1-based and end positions are inclusive, but Python is 0-based and end positions are exclusive.
            # EG GFF 1:10 is 0:10 in Python, where Python returns positions 0 to 9.
            # If gene end is 100, we want GFF positions 48-97.
            # In Python, this is 47-96, but the range is 47:97.
            # This is coding_end-53 to coding_end-3.
            return genome[self.chromosome][self.coding_end-(self.threeprime_homology_length+3):self.coding_end-3].seq
        elif self.strand=='-':
            # If on the reverse strand, the stop codon is at the gene start.
            # Similarly, if gene_start is 1, we want GFF positions 4-53.
            # In Python, this is 3-52, range 3:53, so coding_start+2:coding_start+52.
            return genome[self.chromosome][self.coding_start+2:self.coding_start+(self.threeprime_homology_length+2)].reverse_complement().seq
        else:
            return None

    def __get_primers(self):
        fiveprimers = threeprimers = []
        if self.strand == '+':
            fiveprimers  = get_primers(genome[self.chromosome][self.coding_start-1:self.coding_start+250].seq, self.num_results)
            threeprimers = get_primers(genome[self.chromosome][self.coding_end-250:self.coding_end].seq, self.num_results)
        elif self.strand == '-':
            fiveprimers  = get_primers(genome[self.chromosome][self.coding_end-250:self.coding_end].reverse_complement().seq, self.num_results)
            threeprimers = get_primers(genome[self.chromosome][self.coding_start-1:self.coding_start+250].reverse_complement().seq, self.num_results)
        
        return fiveprimers, threeprimers
        
    def print_plasmid(self, plasmid_type):
        output = f"{self.gene_name}\t{self.chromosome}\t{self.strand}" + \
                 f"\t{self.gene.start}\t{self.gene.end}\t{self.flank_start}\t{self.flank_end}\t{self.coding_start}\t{self.coding_end}"
        
        output += ''.join([f"\t{self.fiveprime_regions[i][1]}\t{GC(self.fiveprime_regions[i][1])}\t{self.fiveprime_regions[i][0]}" for i in range(self.num_results)])
        
        output += f"\t{self.threeprime_region}\t{self.threeprime_gc}"
        
        output += f"\t{self.fiveprimers[0]}\t{self.fiveprimers[1]}\t{self.threeprimers[0]}\t{self.threeprimers[1]}"
        
        if plasmid_type == 'BAC':
            output += ''.join([f"\t{self.bacs[i]}\t{self.bac_wells[i]}" for i in range(self.num_results)])
        elif plasmid_type == 'Fosmid':
            output += ''.join([f"\t{self.fosmids[i]}" for i in range(self.num_results)])

        return output


print(f"Loading genome sequence {args.genomefasta}", file=sys.stderr)
genome = SeqIO.to_dict(SeqIO.parse(gzip.open(args.genomefasta, "rt"), 'fasta'))

wells = {}
print(f"Loading well numbers for BACs", file=sys.stderr)
with open(args.wells, 'r') as w:
    for line in w:
        bac, well = line.strip().split('\t')
        wells[bac] = well

print(f"Loading gene list {args.genelist}", file=sys.stderr)
genes = []
if args.genelist:
    with open(args.genelist, 'r') as gl:
        for line in gl:
            genes.append(line.strip())

genedb, genedb_filename = load_annotation(args.genegff, args.genedatabase)

if not genes:
    for gene in genedb.features_of_type('gene'):
        genes.append(gene.id)

genes = sorted(genes, key=lambda x: (genedb[x].seqid.split('_')[0], int(genedb[x].seqid.split('_')[1]), genedb[x].start))

print("Loading plasmids", file=sys.stderr)
bacs, fosmids = get_plasmid_locations(args.plasmidpairs)

print(f"Processing {len(genes)} genes of interest...", file=sys.stderr)

bacout = open(args.outputstub + ".bacs.tsv", 'w')
fosmidout = open(args.outputstub + ".fosmids.tsv", 'w')

header = "GeneID\tChromosome\tStrand\tGeneStart\tGeneEnd\tFlankStart\tFlankEnd\tCodingStart\tCodingEnd" + \
       ''.join([f"\t5'50bpHomologyRegion{i}\tGC%{i}\t5'UpstreamPosition{i}" for i in range(1,6)]) + \
       "\t3'50bpHomologyRegion\tGC%" + \
       "\t5'primerF1\t5'primerR1\t5'AmpliconSize1\t5'primerF2\t5'primerR2\t5'AmpliconSize2" + \
       "\t3'primerF1\t3'primerR1\t3'AmpliconSize1\t3'primerF2\t3'primerR2\t3'AmpliconSize2"

bac_header    = header + ''.join([f"\tPlasmidID{i}\tPlasmidEnds{i}\tPlasmidStart{i}\tPlasmidEnd{i}\tPlasmidLength{i}\tBACWell{i}" for i in range(1,6)])
fosmid_header = header + ''.join([f"\tPlasmidID{i}\tPlasmidEnds{i}\tPlasmidStart{i}\tPlasmidEnd{i}\tPlasmidLength{i}" for i in range(1,6)])

print(bac_header,    file=bacout)
print(fosmid_header, file=fosmidout)

for gene in genes:
    gene_data = Gene(gene, genedb, bacs, fosmids, wells, args.num_results, \
                     args.fiveprime_flank_length, args.threeprime_flank_length, \
                     args.fiveprime_utr_flank, args.threeprime_utr_flank, \
                     args.fiveprime_homology_length, args.threeprime_homology_length)
                     
    print(gene_data.print_plasmid("BAC"), file=bacout)
    print(gene_data.print_plasmid("Fosmid"), file=fosmidout)

bacout.close()
fosmidout.close()
