## Supplementary material for "A recombineering pipeline to clone large and complex genes in Chlamydomonas": Supplemntal Code: Check_primers-output-sorter.txt

#!/usr/bin/env python3
# -*- coding: utf-8 -*-

"""
CHECK_PRIMERS OUTPUT SORTER
@author Tom Emrich-Mills (2020)
Tested in python3.7

Usage:
    
The Primer3 check_primers module outputs a results summary in Boulder-IO 
format for each primer or pair of primers supplied in the input.
An example output that demonstrates the format for single primers can be seen below.
In this example, the primer was rejected (okay 0) due to breaching a secondary
structure score threshold (high hairpin stability 1).

    SEQUENCE_ID=Sequence1
    SEQUENCE_PRIMER=ATGATGGGCCCCACACCCACTTCATC
    PRIMER_TASK=check_primers
    PRIMER_LEFT_EXPLAIN=considered 1, high hairpin stability 1, ok 0
    PRIMER_LEFT_NUM_RETURNED=0
    PRIMER_RIGHT_NUM_RETURNED=0
    PRIMER_INTERNAL_NUM_RETURNED=0
    PRIMER_PAIR_NUM_RETURNED=0
    =

An example output that demonstrates the format for pairs can be seen below. 
In this example, the forward and reverse primers were rejected (ok 0) due to breaching 
a Tm threshold (tm 1), and the left primer also breached the length threshold (PRIMER_MAX_SIZE).
    
    SEQUENCE_ID=Sequence2vs3
    SEQUENCE_PRIMER=ATGAGCAAAGTTTCAAGGATTCTGCATAGGAAGAG
    SEQUENCE_PRIMER_REVCOMP=CTGTGACATCTGCGCCTGCGAGTC
    PRIMER_TASK=check_primers
    PRIMER_WARNING=Specified left primer > PRIMER_MAX_SIZE
    PRIMER_LEFT_EXPLAIN=considered 1, high tm 1, ok 0
    PRIMER_RIGHT_EXPLAIN=considered 1, high tm 1, ok 0
    PRIMER_PAIR_EXPLAIN=considered 0, ok 0
    PRIMER_LEFT_NUM_RETURNED=0
    PRIMER_RIGHT_NUM_RETURNED=0
    PRIMER_INTERNAL_NUM_RETURNED=0
    PRIMER_PAIR_NUM_RETURNED=0
    =

Check_primers output sorter extracts the reasons for rejection for all rejected 
primers (and pairs if available) and outputs a CSV file, enabling fast quantification 
of the number and type of any warnings or reasons for rejection produced by the module. 
The script can be run separately depending on whether the user wishes to analyse pairs 
of primers against each other or just a list of single primers - simply specify the use
of lines 105-110 or lines 115-121.  
The script is run from within a python development environment.
Add your Boulder-IO file produced by Primer3 in place of ./p3output.txt.
Add an output file address in place of ./p3output_sorted.csv.

"""

import csv

input_boulder_loc = "./p3output.txt"
output_csv_loc = "./p3output_sorted.csv"

names_list = []
F_list = []
R_list = []
Warning_list = []
Left_explain = []
Right_explain = []
Pair_explain = []
F_okay = []
R_okay = []
P_okay = []

with open(input_boulder_loc, 'r') as f:
    reader=csv.reader(f)
    for row in reader:
        if row[0].startswith("SEQUENCE_ID"):
            names_list.append(str(row)[14:-2])
        if row[0].startswith("SEQUENCE_PRIMER="):
            F_list.append(str(row)[18:-2])
        if row[0].startswith("SEQUENCE_PRIMER_R"):
            R_list.append(str(row)[26:-2])
        if row[0].startswith("PRIMER_WARNING"):
            char_loc = str(row).find(">")
            Warning_list.append(str(row)[char_loc+2:-2])
        else:
            Warning_list.append("")
        if row[0].startswith("PRIMER_LEFT_EXPLAIN"):
            F_okay.append(str(row)[-3:-2])
            Left_explain.append(str(row)[39:-13])
        if row[0].startswith("PRIMER_RIGHT_EXPLAIN"):
            R_okay.append(str(row)[-3:-2])
            Right_explain.append(str(row)[40:-13])
        if row[0].startswith("PRIMER_PAIR_EXPLAIN"):
            high_loc = str(row).find("high")
            if str(row)[high_loc:79] == "high end":
                Pair_explain.append(str(row)[high_loc:85])
            else:
                Pair_explain.append("")
            
f.close()

#"""
### For single primers ###
    
with open(output_csv_loc, 'w') as g:
    writer=csv.writer(g)
    writer.writerow(['Name', 'Seq', 'Okay', 'Reason'])
    for i in range(len(names_list)):
        current_row = [names_list[i]] + [F_list[i]] + [F_okay[i]] + [Left_explain[i]]
        writer.writerow(current_row)

"""
### For pairs ###

with open(output_csv_loc, 'w') as g:
    writer=csv.writer(g)
    writer.writerow(['Name', 'Seq F', 'F okay', 'F reason', 'Seq R', 'R okay', 'R reason', 'Pair reason'])
    for i in range(len(names_list)):
        current_row = [names_list[i]] + [F_list[i]] + [F_okay[i]] + [Left_explain[i]] \
            + [R_list[i]] + [R_okay[i]] + [Right_explain[i]] + [Pair_explain[i]]
        writer.writerow(current_row)
"""

g.close()
