## Supplementary material for "A recombineering pipeline to clone large and complex genes in Chlamydomonas": Supplemntal Code: README.txt

Supplemental Code

Developed by Tom Emrich-Mills and John W. Davey

Associated manuscript: A recombineering pipeline to clone large and complex genes in Chlamydomonas, by Emrich-Mills et al. (2020)

All files are available in the associated GitHub repository: https://github.com/TZEmrichMills/Chlamydomonas_recombineering


1. BACSearcher.txt

The BACSearcher python script, including brief usage instructions.
For more detailed information regarding function, usage and precautions, see Supplemental Method 2.


2. BACs_fosmids.pairs.tsv

Precursor file required by BACSearcher to predict BAC coverage for each gene.
For usage instructions, see Supplemental Method 2A; this file corresponds to precursor file I.
The file contains start and end coordinates for all valid BACs and fosmids within the Chlamydomonas BAC library, as well as the lengths of each. 
BACs are identified by a PTQ number and fosmids are identified by a VTP number. 
Plasmids are defined as valid if their start and end sequences are mapped to the same chromosome, with one end mapped to the sense strand and the other to the anti-sense strand. 
These criteria exclude 3179 of the 11,676 BACs (27.2%) and 20,696 of the 56,276 fosmids (36.8%) that are annotated against the Chlamydomonas genome (v5.5). 
Due to the way the BACs have been annotated against the genome, some valid plasmids have more than two ends mapped to the same chromosome and so are represented more than once (see Supplemental Method 2B for additional information).


3. BAC_wells.txt

Precursor file required by BACSearcher in order to provide plate coordinates for any predicted BACs within the Chlamydomonas BAC library.
For usage instructions, see Supplemental Method 2A; this file corresponds to precursor file V. 


4. TRF-output-sorter.txt

Python script intended for processing the output information produced by Tandem Repeats Finder (Benson, 1999) into CSV format. 
Usage instructions are included at the top of the script and in Supplemental Method 4A.
See Supplemental Method 3A for details on Tandem Repeats Finder usage.


5. WindowMasker-output-sorter.txt

Python script intended for counting the number of repeat sequences masked by WindowMasker (Morgulis et al., 2006).
The script takes the interval TXT file produced by WindowMasker and outputs a CSV file containing the number or repeats per input sequence.
Usage instructions are included at the top of the script and in Supplemental Method 4B.
See Supplemental Method 3C for details on WindowMasker and its usage.


6. Check_primers-output-sorter.txt

Python script intended to enable quick analysis of the output from the Primer3 check_primers module (Rozen and Skaletsky, 2000). 
Usage instructions are included at the top of the script and in Supplemental Method 4C. 
See Supplemental Method 3D for details of the Primer3 check_primers module.

-
