## Supplementary material for "A recombineering pipeline to clone large and complex genes in Chlamydomonas": Supplemntal Code: TRF-output-sorter.txt

#!/usr/bin/env python3
# -*- coding: utf-8 -*-

"""
TRF OUTPUT SORTER
@author Tom Emrich-Mills and G.E.Mayneord (2020)
Tested in python3.7

Usage:

Tandem Repeats Finder (Benson, 1999) outputs a data file containing lists of repeats 
for every entry (e.g., gene sequence supplied in fasta format). An example output that
demonstrates the format of this data file can be seen below. 
In this example, Sequence1 was found to contain 9 repeats, each with accompanying
analysis information (see https://tandem.bu.edu/trf/trf.definitions.html for details).

    Sequence: Sequence1
    
    Parameters: 2 5 7 80 10 20 2000
    
    257 270 5 2.8 5 100 0 28 0 42 42 14 1.45 CGGCT CGGCTCGGCTCGGC
    440 449 4 2.5 4 100 0 20 0 80 20 0 0.72 CCCG CCCGCCCGCC
    1329 1346 8 2.3 8 100 0 36 22 66 11 0 1.22 CCCGCAAC CCCGCAACCCCGCAACCC
    1529 1549 11 1.9 11 90 0 35 14 14 61 9 1.55 GTGAAGGCGGC GTGGAGGCGGCGTGAAGGCGG
    1531 1569 17 2.4 16 86 4 55 17 12 64 5 1.46 GGAGGAGGCGTGAGGC GGAGGCGGCGTGAAGGCGGAGGAGGCGTGAGGCGGGGGA
    1565 1585 6 3.3 6 86 6 26 4 4 66 23 1.30 GGGGTT GGGGATGGGGTTGGGGCTTGG
    1656 1666 4 2.8 4 100 0 22 27 27 45 0 1.54 GCAG GCAGGCAGGCA
    1777 1789 6 2.2 6 100 0 26 15 15 46 23 1.83 TAGGGC TAGGGCTAGGGCT
    1855 1869 4 3.8 4 90 0 23 26 13 33 26 1.93 ATGC ATGCATGCATGGATG

TRF output sorter reads the data file produced by Tandem Repeats Finder and reformats the information 
into a CSV file in which each reported repeat sequence (and its accompanying analysis information) 
is present on its own row, preceded by the name of the gene in which it was detected. 
This enables straightforward analysis of the number, length, type and identity of tandem 
repeats per gene, as well as the number of genes containing at least one tandem repeat. 
The script is run from within a python development environment, and is currently set to 
filter out any repeats that do not show >=90% sequence identity (line 115).
Replace 'Path\Sequences.fa.2.5.7.80.10.20.2000.dat' with your data file address.
Replace 'Path\Sequences_sorted' with an output file location below. 

"""

import csv
import os
from tqdm import tqdm

#input_data_file_address = "./Sequences.fa.2.5.7.80.10.20.2000.dat"
#output_file_loc = "./Sequences_sorted.csv"

input_data_file_address = "/Users/Tommy/Dropbox/Tommy/02 Previous labs/Mackinder Lab/Recombineering manuscript/Figures/Genome analysis - Repeats/190426_TRF/Example.fa.2.5.7.80.10.20.2000.dat"
output_file_loc = "/Users/Tommy/Dropbox/Tommy/02 Previous labs/Mackinder Lab/Recombineering manuscript/Figures/Genome analysis - Repeats/190426_TRF/Example.fa.2.5.7.80.10.20.2000.90.csv"

with open(input_data_file_address) as e:
   lines = e.readlines()
   newLines = []
   for line in lines:
      newLine = line.strip().split()
      newLines.append( newLine )

with open('output1.csv', 'w', newline='') as f:
   file_writer = csv.writer(f)
   file_writer.writerows( newLines )

e.close()

with open('output1.csv') as input, open('output2.csv', 'w', newline='') as output:
    non_blank = (line for line in input if line.strip())
    output.writelines(non_blank)

f.close()

total_data_stored=[]
current_list_stored=[]

with open('output2.csv') as g:
    reader=csv.reader(g)
    for row in reader:
        if row[0] == 'Sequence:':
            total_data_stored.append(current_list_stored)
            current_list_stored=[]
            current_list_stored.append(row[1])   
        else:
            current_list_stored.append(row)

total_data_stored.append(current_list_stored)
# delete all the data before the first time the word 'Sequence' appears
del total_data_stored[0] 
Parameters_list = total_data_stored[0][1]
for i in range(len(total_data_stored)):
        total_data_stored[i].remove(total_data_stored[i][1])
        
for i in range(len(total_data_stored)):      
    if len(total_data_stored[i]) == 1:
        if_no_reps = ['No repeats detected']
        total_data_stored[i] = total_data_stored[i] + [if_no_reps]

with open(output_file_loc, 'w', newline='') as h:
    writer=csv.writer(h)
    writer.writerow(['Gene', 'Result #', 'From pos', 'To pos', 'Period size', 'Copy number', 'Rep region length',
                     'Consensus size','Percent matches', 'Percent indels', 'Score',
                     'A', 'C', 'G','T', '%GC', 'Entropy', 'Repeat motif', 'Repeat region'])

    for i in tqdm(range(len(total_data_stored))):
        current_header=total_data_stored[i][0]
        for j in range(1, (len(total_data_stored[i])), 1):
            current_list=total_data_stored[i][j]
            current_list.insert(0, j)
            current_list=current_list[0:len(current_list)] 
            current_list=[current_header] + current_list
            if current_list[2] == 'No repeats detected':
                extra_columns=['', '', '', '', '', '', '', '', '', '', '', '', '', '', '', '']
                current_list = current_list[0:3] + extra_columns
                writer.writerow(current_list)
            else:
                if float(current_list[7])>=90: 
                    RepRegionLength = len(current_list[16])
                    GC_content = (str(current_list[16]).count("C") + str(current_list[16]).count("G"))/RepRegionLength
                    Rounded_GC_content = round(float(GC_content), 2)
                    current_list = current_list[0:6] + [RepRegionLength] + current_list[6:14] + [Rounded_GC_content] + current_list[14:]     
                    writer.writerow(current_list)  

os.remove('output1.csv')
os.remove('output2.csv')        
h.close()

print('==========')
print ('Finished')
