## Supplementary material for "A recombineering pipeline to clone large and complex genes in Chlamydomonas": Supplemntal Code: WindowMasker-output-sorter.txt

#!/usr/bin/env python3
# -*- coding: utf-8 -*-

"""
WINDOWMASKER OUTPUT SORTER
@author Tom Emrich-Mills (2020)
Tested in python3.7

Usage:

WindowMasker (Morgulis et al., 2006) accepts one or multiple nucleotide 
sequences and outputs an intervals file containing the name of each analysed 
sequence followed by the start and end positions of each masked repetitive 
sequence found within the entry (one run per line). 
If a FASTA file is supplied, the WindowMasker output file has the following format:

    >Sequence1
    824 - 835
    1175 - 1186
    1210 - 1370
    >Sequence2
    377 - 388
    465 - 479
    1437 - 1455
    1705 - 1716

WindowMasker output sorter uses the python data analysis library (pandas)
to simply count the number of intervals per entry sequence, equating to the 
number of masked repeats per sequence. The script outputs a CSV file to enable 
the quick generation of values for the average number of masked regions per 
sequence and number of sequences with at least one masked region. 
The script is run from within a python development environment.
Add your interval file address in  place of ./WMoutput.txt, below.
Add an output file location to in place of ./WMoutput_sorted.csv.

"""

import pandas as pd

input_interval_file_address = "./WMoutput.txt"
output_csv_location = "./WMoutput_sorted.csv"

with open(input_interval_file_address, 'r') as g:                                                   
    data = pd.read_csv(g, names=['Content']) 
    index_list = []
    content_list = []
    differences_list = []
    df=pd.DataFrame(data)
    # Temporarily add an end row to aid with downstream counting
    end_row = pd.DataFrame([['>End']], columns=['Content'])
    df1=df.append(end_row, ignore_index=True)
    only_genes = df1[df1.Content.str.get(0).isin(['>'])]
    with_index = only_genes.reset_index()
    with_index['Content'] = with_index['Content'].str[1:]
    only_index = with_index[['index']].copy()
    # Define a new df containing the differences between the repeat intervals
    differences = only_index.diff(axis = 0, periods = -1) 
    with_index['Reps per gene'] = differences
    with_index['Reps per gene'] = with_index['Reps per gene'].apply(str).str[1:]
    del with_index['index']
    full_table = with_index.iloc[:-1]
full_table.to_csv(output_csv_location, index=False)

g.close()
