## Supplemental Data for "A recombineering pipeline to clone large and complex genes in Chlamydomonas"

#### Supplemental Figure 1. Plasmid map for recombineering vectors

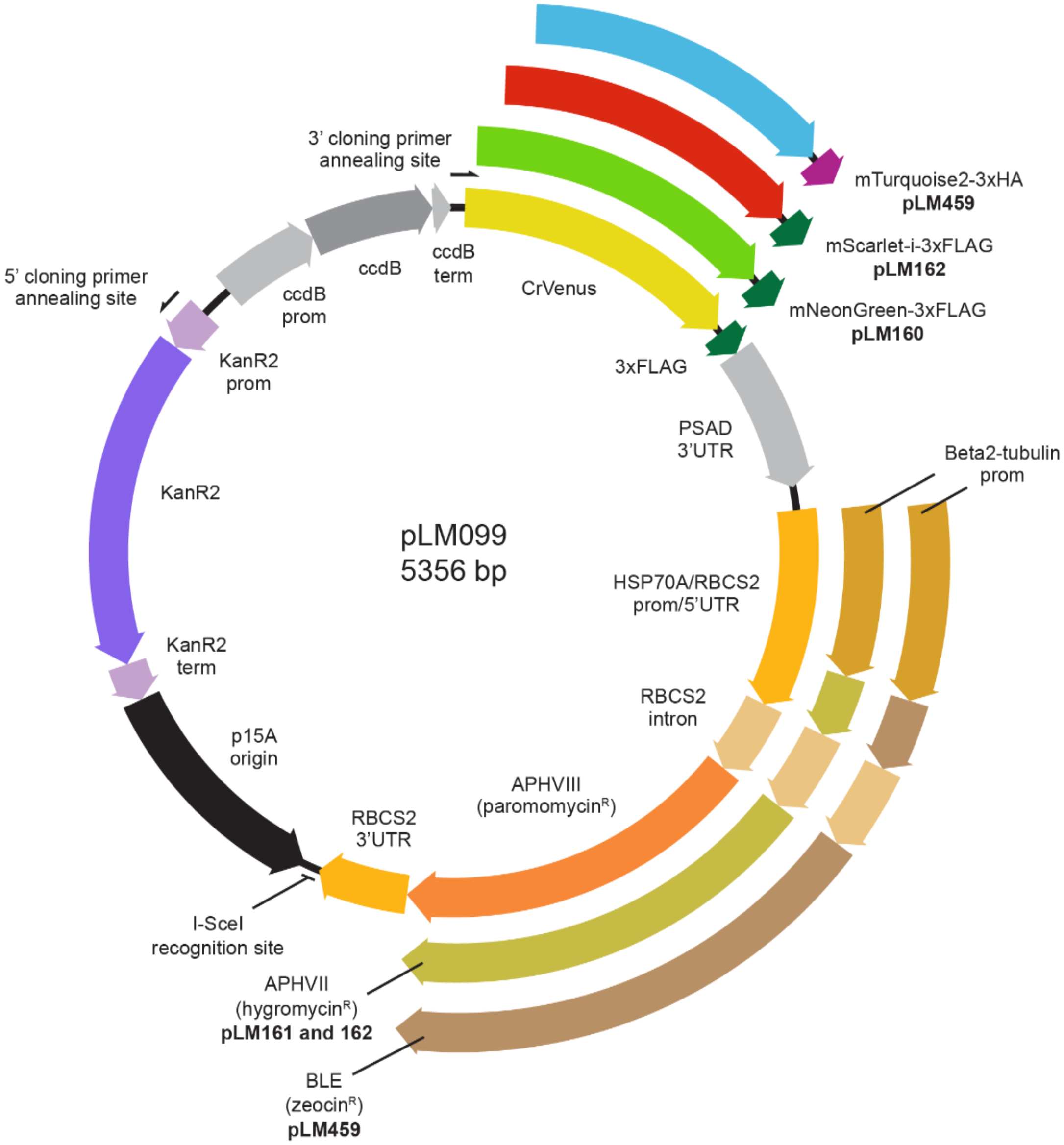

**Supplemental Figure 1** Plasmid map for pLM099 and derivative recombineering vectors.
PCR amplification with 5' and 3' cloning primers at the annealing sites shown results in a ~4.6 kbp linear cassette for recombineering target genes in-frame with a fluorescent protein and affinity tag. For each recombineering vector, the fluorescent protein sequence is preceded by a short, flexible linker (GGLGGSGGR) and followed by a tri-glycine linker prior to the affinity tag. All four fluorescent protein-affinity tag cassettes are terminated with the PSAD 3'UTR. All three Chlamydomonas selection cassettes (paromomycin$^R$, hygromycin$^R$ and zeocin$^R$) are terminated with the RBCS2 3'UTR. The same RBCS2 intron is present in all three Chlamydomonas selection cassettes to improve expression but is only inter-exonic in the hygromycin and zeocin resistance cassettes.

### Supplemental Figure 2. Batch-scale recombineering results

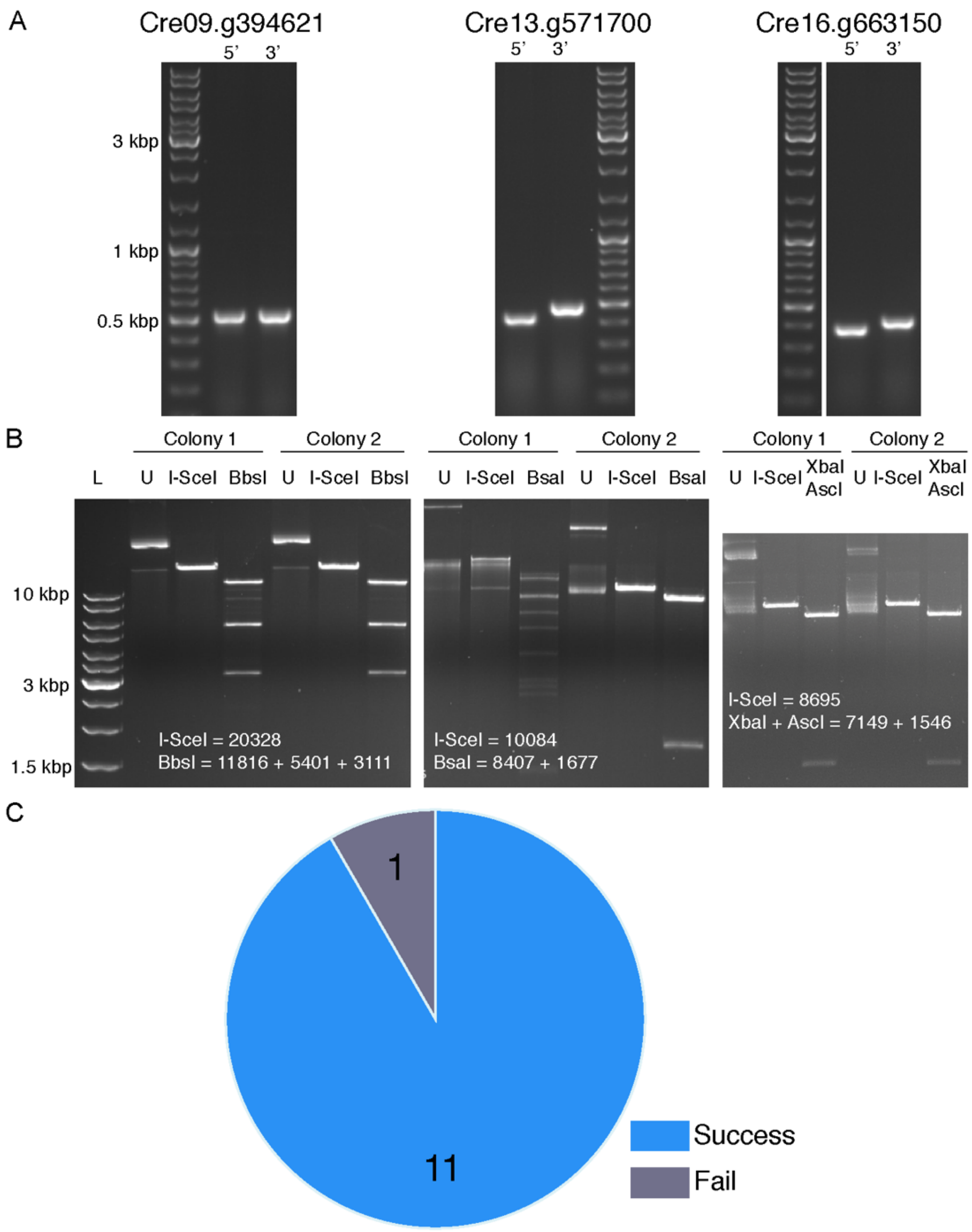

**Supplemental Figure 2** Batch-scale recombineering results.
**A** Three examples of colony PCRs to check for presence of target genes in BACs. Primer pairs were designed to the 5' and 3' end of each target gene. All amplicons were of the expected size.
**B** Restriction digest checks for isolated recombineered plasmids from two colonies per gene, corresponding to the same genes as in **A**. Expected sizes are shown in bp. Note that colony 1 for Cre13.g571700 gives the incorrect size and banding patterns after digestion indicating incorrect recombination. U: undigested.
**C** Overall batch-scale recombineering success for 12 target genes.

#### Supplemental Method 1. Protocols for batch- and large-scale recombineering

**Background**

The recombineering pipelines detailed in these protocols enable the one-step sub-cloning and tagging of target regions of Chlamydomonas gDNA at the C-terminus with the fluorescent proteins CrVenus (pLM099 and pLM161), mNeonGreen (pLM160), mScarlet-i (pLM162) or mTurquoise2 (pLM459). Sub-cloning occurs by homologous recombination between the target gDNA and a recombineering cassette, resulting in a recombineering plasmid product containing the target gene/region in-frame with a fluorescent protein and affinity tag. Target gDNA is contained in bacterial artificial chromosomes (BACs) from the Chlamydomonas BAC library (*E. coli* strain DH10B). Each clone in the library contains a single copy of the respective BAC, consisting of a stretch of Chlamydomonas gDNA and a backbone derived from pBeloBAC11, which contains a chloramphenicol resistance gene. The library is available from the Chlamydomonas Resource Centre (https://www.chlamycollection.org/).

**Primer design & PCR**

Each recombineering reaction requires a target-specific recombineering cassette, amplified by PCR from either pLM099 or its derivate vectors pLM160, 161, 162 or 459 (see Figure 5 and Supplemental Figure 1). The same forward (26 bp) and reverse (19 bp) primer annealing sequences can be used to amplify all vectors (see 1C., appendix). The forward primer annealing sequence for each reaction is preceded by a target-specific 50 bp homology arm corresponding to a site at least 2000 bp upstream of the annotated ATG of a target gene contained within the BAC. The reverse primer annealing sequence in each reaction is preceded by a similar homology region corresponding to the 50 bp immediately upstream of the target gene stop codon. Example primers containing homology arms can be seen in the appendix. Five suitable homology regions for almost every gene in the nuclear genome are provided in Supplemental Data Set 1. The PCR product resulting from the amplification of pLM099 or its derivative vectors is referred to as the recombineering cassette in the following protocols. See Supplemental Method 2A for instructions to generate primer homology arms to produce larger or smaller upstream flanking regions.

#### A. Batch-scale recombineering pipeline protocol

##### Overview

The protocol takes place over 5 days; Day 0 is a preparation day, the pipeline takes 4 days to complete. See Figure 2 for a graphical summary of the procedure. Here the protocol is optimized for low sample numbers using 50 ml culturing flasks and 15 ml growth tubes. For a high throughput protocol using volumes suitable for multi-well plate-scale cloning, see next section.

**Materials**

Antibiotics (working concentrations):
- Chloramphenicol 12.5 µg/ml ($chl_{12.5}$)
- Tetracycline 5 µg/ml ($tet_5$)
- Kanamycin 25 µg/ml ($kan_{25}$)

Media (sterile):
- Lysogeny broth (LB)
- Yeast extract nutrient broth (YENB)
- Super optimal broth with catabolite repression (SOC)

10% glycerol (sterile)
10% L-arabinose (sterile)
Nuclease-free water
Oligonucleotides (see Supplemental Data Set 3)
Plasmid extraction kit
Gel extraction/PCR purification kit
High fidelity polymerase
I-SceI homing endonuclease
Other common restriction enzymes (gene dependant)

**Equipment**

Incubation room/chamber for 30°C and 37°C, with orbital shaking equipment
Spectrophotometer for measuring bacterial OD
Nanodrop spectrophotometer
Electroporation cuvettes (2 mm gap)
Electroporator capable of 2500 V pulses (e.g. GenePulser II, Bio-Rad)
Refrigerated microcentrifuge
PCR thermocycler
2 ml Eppendorf tubes
15 ml bacterial culturing tubes
50 ml conical flask
90 mm petri dish

**Summary**

Day 0. Inoculation of liquid culture with bacteria containing the target BAC. Extraction of plasmid pRed and plasmid pLM099.* Design of cloning primer homology arms.* Amplification of target-specific recombineering cassette from pLM099.

Day 1. Electrotransformation of BAC-containing clones with pRed. Outgrowth and selection with chloramphenicol and tetracycline.

Day 2. Induction of the Redαβγ and recA genes with L-arabinose. Electrotransformation with target-specific recombineering cassette. Outgrowth and selection with kanamycin.

Day 3. Selection of positive colonies.

Day 4. Plasmid product extraction and restriction digest analysis.

* Any time before the procedure.

**Procedure**

Day 0:

Pre-prepared: pRed plasmid, purified recombineering cassette (see appendix for PCR protocol).

*It is best to purify the recombineering cassette PCR product as close to the day of use as possible, with no freeze thaw cycles in between.*

1. Using a single colony of DH10B *E. coli* containing the BAC of interest, inoculate a 50 ml flask containing 20 ml of YENB with $chl_{12.5}$ and grow overnight at 37°C with shaking at 200-220 rpm.

Day 1:

1. In a 15 ml culturing tube, inoculate 4 ml of YENB containing $chl_{12.5}$ with 120 μl of the overnight BAC clone culture and grow for 2 h at 37°C with vigorous shaking until it reaches $OD_{600}$ of >2.

Pre-cool microcentrifuge to 4°C.
Dilute pRed plasmid to 0.1 ng/μl in nuclease-free water (100 μl per reaction).
Keep the following on ice: 10% glycerol, electroporation cuvettes (1 per sample), 2 ml tubes and the diluted pRed.
Add 800 μl of SOC to 2 ml tubes and keep at room temperature.

2. Transfer 2 ml of the saturated overnight growth of the BAC strain to a pre-cooled 2 ml tube and incubate on ice for 2 min.
3. Spin culture in a pre-cooled 4°C microcentrifuge for 10 min at 5000 x g.
4. Pour off supernatant and tap tube to remove residual supernatant before placing back on ice.
5. Resuspend pellet in 1 ml of 10% glycerol by gently pipetting up and down.
6. Centrifuge at 5000 x g for 10 min at 4°C.
7. Carefully remove all supernatant (glycerol-washed pellet is much looser) and place back on ice.
8. Resuspend in 100 μl of 0.1 ng/μl of pRed plasmid and transfer to a pre-chilled cuvette.
9. Dry cuvette and electroporate at 2500 V, 400 Ω and 25 μF.*
10. Immediately transfer cells into a 2 ml tube containing 800 μl of SOC.
11. Allow 90 min of recovery growth at **30°C** with shaking at 200-220 rpm.

12. Inoculate 20 ml of fresh YENB containing $chl_{12.5}$ & $tet_5$ with 400 µl of outgrowth culture. Grow in a 50 ml flask overnight for at least 16 h at 30°C with vigorous shaking.

*To check the efficiency of this step, the remaining outgrowth can be plated onto LB-agar with $chl_{12.5}$ & $tet_5$ and incubated at 30°C overnight. The presence of multiple colonies indicates success, though colony number can vary from ~30 to >500 depending on the BAC clone.*

* These settings were optimised for a Bio-Rad GenePulser II and may vary for other electroporators.

Day 2:

1. Inoculate 4 ml of fresh YENB containing $chl_{12.5}$ & $tet_5$ with 600 µl of the overnight growth. Incubate at **30°C** with vigorous shaking for ~3 h (or until $OD_{600}$ >2)

*It is important to achieve a saturated growth at this stage – this step can be extended to 4 h if needed.*

Pre-cool microcentrifuge to 4°C.
Dilute target-specific recombineering cassette to 5 ng/µl in nuclease-free water (100 µl per reaction).
Keep the following on ice: 10% glycerol, electroporation cuvettes (1 per sample), 2 ml tubes, the diluted target-specific recombineering cassette (5 ng/µl) and 10% L-Arabinose (80 µl per sample).
Add 800 µl of SOC to a 2 ml tubes and keep at room temperature.

2. Add 80 µl of 10% L-arabinose to the culture and incubate at **37°C** for 1 h with shaking.
3. Transfer 2 ml to a pre-cooled 2 ml tube and incubate on ice for 2 min.
4. Spin culture in a precooled 4°C microcentrifuge for 10 min at 5000 x g.
5. Pour off supernatant and tap tube to remove residual supernatant before placing back on ice.
6. Resuspend pellet in 1 ml of 10% glycerol by gently pipetting up and down.
7. Centrifuge at 5000 x g for 10 min at 4°C.
8. Carefully remove all supernatant and place back on ice.
9. Resuspend in 100 µl of recombineering cassette (5 ng/µl) and transfer to a pre-chilled cuvette.
10. Dry cuvette and electroporate at 2500 V, 400 Ω and 25 µF.
11. Immediately transfer cells into a 2 ml tube containing 800 µl of SOC.
12. Allow 90 min of recovery growth at 37°C with shaking.
13. Plate all outgrowth onto 2 x LB-agar plates containing $kan_{25}$ (~450ul on each), air dry, and incubate plates at 37°C overnight (at least 16 h).

Day 3:

1. Choose 3 or more colonies and inoculate each into 4 ml of fresh LB containing $kan_{25}$ and grow at 37°C overnight with shaking. For larger targets, this can be increased to 6 ml.

Day 4:

1. Extract plasmids, elute in 30 µl nuclease-free water or EB and measure the concentration on a nanodrop spectrophotometer.
2. Set up two 200-500 ng restriction digestion reactions (one single-cutter and one multi-cutter). Analyse on an agarose gel.

*From 4 ml of growth, the yield is typically 40-120 ng/µl of plasmid product, with an average of 80 ng/µl for target regions ~7 kbp in length. For very large targets increase growth volume to maximise yield.*

#### B. Large-scale recombineering pipeline protocol

##### Overview

The protocol takes place over 5 days; Day 0 is a preparation day, the pipeline takes 4 days to complete. See Figure 2 for a graphical summary of the procedure. Here the protocol is optimized for high-throughput cloning using 96-well plates. Volumes given here are for individual wells or individual 2 ml tubes for some steps. For small-scale batch optimised protocol, see above.

##### Materials

Antibiotics (working concentrations):
- Kanamycin 25 µg/ml ($kan_{25}$),
- Tetracycline 5 µg/ml ($tet_{5}$),
- Chloramphenicol 12.5 µg/ml ($chl_{12.5}$)

Media (sterile):
- Lysogeny broth (LB)
- Yeast extract nutrient broth (YENB)
- Super optimal broth with catabolite repression (SOC)

10% glycerol (sterile)
10% L-arabinose (sterile)
Nuclease-free water
Oligonucleotides (see Supplemental Data Set 3)
96-well plasmid extraction kit
96-well gel extraction/PCR purification kit
High fidelity polymerase
I-SceI homing endonuclease
Other common restriction enzymes (gene dependant)

##### Equipment

Incubation room/chamber for 30°C and 37°C, with orbital shaking equipment
Nanodrop spectrophotometer
2 mm gap electroporation cuvettes or 96-well electroporation plates
Electroporator capable of 2500 V pulses (e.g. GenePulser II, Bio-Rad)
Refrigerated centrifuge with plate adapters
PCR thermocycler with 96-well adapter
12-channel 1 ml pipette
1.5 ml and 2 ml Eppendorf tubes
96-well 2.2 ml deep-well culturing plates
96-well 2 ml square-well culturing plates
96-well PCR plates
90 mm petri dish

**Summary**

Day 0. Inoculation of liquid culture with bacteria containing the target BAC. Extraction of plasmid pRed and plasmid pLM099.* Design of cloning primer homology arms.* Amplification of target-specific recombineering cassette from pLM099.

Day 1. Electrotransformation of BAC-containing clones with pRed. Outgrowth and selection with chloramphenicol and tetracycline.

Day 2. Induction of the Redαβγ and recA genes with L-arabinose. Electrotransformation with target-specific recombineering cassette. Outgrowth and selection with kanamycin.

Day 3. Selection of positive colonies.

Day 4. Plasmid product extraction and restriction digest analysis.

* Prepare before beginning procedure

**Procedure**

Day 0:

Pre-prepared: pRed plasmid, purified recombineering cassette (see appendix for PCR protocol).

*It is best to purify the recombineering cassette PCR product as close to the day of use as possible, with no freeze thaw cycles in between.*

1. Using a single colony of DH10B *E. coli* containing the BAC of interest, inoculate each well of a 2.2 ml deep-well culturing plates containing 1 ml of fresh YENB with $chl_{12.5}$, and grow overnight at 37°C on a plate shaker at 900 rpm.

Day 1:

1. Inoculate 900 µl of fresh YENB containing $chl_{12.5}$ with 40 µl of the overnight BAC clone culture and grow for 2 h at 37°C with shaking at 900 rpm.

Pre-cool centrifuge with plate adapters to 4°C.
Dilute pRed plasmid to 0.1 ng/µl in nuclease-free water (100 µl per reaction).
Keep the following on ice: 10% glycerol (1 ml per sample), 1 x 96-well deep-well culture plate (or 96 x 2 ml tubes), 96 x electroporation cuvettes (1 per sample), and the diluted pRed.
Add 800 µl of SOC per well to an additional 96-well deep-well culturing plate and keep at room temperature.

2. For each well, add the whole saturated overnight BAC clone culture to the corresponding wells in the pre-cooled 96-well plate (or a 2 ml tube), and incubate on ice for 2-5 min.
3. Spin cultures in a pre-cooled 4°C centrifuge for 10 min at 5000 x g.
4. Pour off supernatants and tap inverted plates to remove residual supernatant before placing back on ice.
5. Resuspend pellets in 1 ml of 10% glycerol, gently pipetting up and down.
6. Centrifuge at 5000 x g for 10 min at 4°C.
7. Keeping the plates on ice, carefully remove all supernatant from each well using a 12-channel pipette (glycerol-washed pellets are much looser).

8. Resuspend pellets in 100 µl of 0.1 ng/µl of pRed plasmid and transfer to pre-chilled cuvettes.*
9. Dry each cuvette and electroporate at 2500 V, 400 Ω and 25 µF.**
10. Immediately transfer cells into the corresponding well of the 96-well deep-well culturing plate containing 800 µl of SOC.
11. Allow 90 min of recovery growth at 30°C with shaking.
12. Inoculate 800 µl of fresh YENB containing $chl_{12.5}$ & $tet_5$ with 100 µl of outgrowth culture. Grow overnight for at least 16 h at **30°C** on a plate shaker at 900 rpm.

*To check the efficiency of this step, the remaining outgrowth can be plated onto LB-agar with* $chl_{12.5}$ *&* $tet_5$ *and incubated at 30°C overnight. The presence of multiple colonies indicates success, though colony number can vary from ~30 to >500 depending on the BAC clone.*

* Although not tested, 96-well electroporation plates could be used here instead of individual cuvettes.
** These settings were optimised for a Bio-Rad GenePulser II and may vary for other electroporators.

Day 2:

1. Inoculate 850 µl of fresh YENB containing $chl_{12.5}$ & $tet_5$ with 120 µl of the overnight growth. Incubate at **30°C** with vigorous shaking for ~3 h.

*Optimizing bacterial growth at this stage is key to efficient electroporation. This step can be extended if some wells appear less saturated.*

Dilute target-specific recombineering cassettes to 5 ng/µl in nuclease-free water (100 µl per reaction).
Pre-cool centrifuge with plate adapters to 4°C.
Keep the following on ice: 10% glycerol (1 ml per sample), 1 x 96-well deep-well culture plate (or 96 x 2 ml tubes), 96 x electroporation cuvettes (1 per sample), the diluted target-specific recombineering cassette and 10% L-arabinose (20 µl per sample).
Add 800 µl of SOC to each well of a 96 well 2ml deep culture plate and keep at room temperature.

2. Add 20 µl of 10% L-arabinose to each well and incubate at **37°C** for 1 h with shaking.
3. For each well, add the whole saturated culture to the corresponding wells in the pre-cooled 96-well plate (or to a 2 ml tube), and incubate on ice for 2 min.
4. Spin cultures in a precooled 4°C microcentrifuge for 10 min at 5000 x g.
5. Pour off supernatants and tap inverted plates to remove residual supernatant before placing back on ice.
6. Resuspend pellets in 1 ml of 10% glycerol, gently pipetting up and down.
7. Centrifuge at 5000 x g for 10 min at 4°C.
8. Carefully remove all supernatant and place back on ice.
9. Resuspend in 100 µl of recombineering cassette (5 ng/µl) and transfer to pre-chilled cuvettes.
10. Dry each cuvette and electroporate at 2500 V, 400 Ω and 25 µF.
11. Immediately transfer cells into the corresponding well of the 96-well deep-well culturing plate containing 800 µl of SOC.
12. Allow 90 min of recovery growth at 37°C with shaking.
13. Plate all outgrowth onto 2 x LB-agar plates containing $kan_{25}$ (~450ul on each), air dry, and incubate plates at 37°C overnight (at least 16 h).

Day 3:

1. Pick 1 colony per sample and inoculate into a 96-well 2 ml square-well culturing plate containing 1.2 ml of fresh LB with $kan_{25}$ and grow at 37°C overnight with vigorous shaking.

*Well shape significantly affects growth at this stage. We used the recommended square-well culturing plates supplied with the Wizard 96 plasmid purification kit (Promega).*

Day 4:

1. Using a 96-well plasmid isolation kit, extract plasmids, elute in nuclease-free water and measure the concentrations using a nanodrop spectrophotometer.
2. Set up two 100-200 ng restriction digestion reactions per sample in 96-well PCR plates (one single-cutter and one multi-cutter). Analyse on agarose gels.

*From 1.2 ml of growth, the yield is typically 15-50 ng/µl of plasmid product, with an average of 30 ng/µl for target regions ~7 kbp in length. For very large targets the yield may be lower*.

#### C. Appendix

##### PCR protocols for amplifying target-specific recombineering cassette from pLM099

Standard protocol - 50 µl reaction:

| | |
|---|---|
| Nuclease-free water | 28.1 µl |
| NEB GC buffer (5X) | 10 µl |
| DMSO (100%) | 4.5 µl |
| pLM099 (10 ng/µl) | 1 µl |
| dNTPs (10 mM) | 1 µl |
| Primers (10 µM) | 2.5 µl each |
| Phusion HS II | 0.4 µl |

Betaine-enhanced protocol - 50 µl reaction:

| | |
|---|---|
| Nuclease-free water | 19.6 µl |
| NEB GC buffer (5X) | 10 µl |
| DMSO (100%) | 3 µl |
| Betaine (5 M) | 10 µl |
| pLM099 (10 ng/µl) | 1 µl |
| dNTPs (10 mM) | 1 µl |
| Primers (10 µM) | 2.5 µl each |
| Phusion HS II | 0.4 µl |

Thermocycler setup:

| | | |
|---|---|---|
| 98°C | 1 min | |
| 98°C | 30 sec | x 35 |
| 63°C | 30 sec | x 35 |
| 72°C | 100 sec | x 35 |
| 72°C | 10 min | |

Check 1 µl on an agarose gel – the PCR product should be ~4600 bp. Purify the PCR product using a PCR purification protocol. Elute with water. Dilute with water.

The inclusion of 9% DMSO helps disrupt secondary structure formation of the high-GC content cassette. The inclusion of 6% DMSO plus 1 M betaine can yield success for reactions that prove difficult using the standard protocol. Phusion Hot Start II polymerase was obtained from ThermoFisher Scientific. 5X Phusion GC buffer was obtained from New England Biolabs. The same protocol can be used to amplify recombineering cassettes from pLM160, 161, 162 and 459.

##### Cloning primer sequences examples

5' target-specific homology arms (>2000 bp upstream of the gene start codon)
3' target-specific homology arms (immediately upstream of the gene stop codon)
Annealing sequence common to all recombineering plasmids

Cre09.g394621
5': AAAGTAATTACAATATCCCTTGATATAAACTACAGTGGTATATCTAAGCCGAAGATCCTTTGATCTTTTCTACGGG
3': CCAGTCCCGTGCGCCGCACCGCTATCCCGGCCAACTGGCGTGACGCACTGGGAGATCTGGGTGGCTCCG

Cre06.g273050
5': TCCTCACATTTACTTCTTCTTGGCGACACTGTGGATGGTTTACAGCGTGCGAAGATCCTTTGATCTTTTCTACGGG
3': AATACCGCCGGAGAGCGCCCGCCAAGCCGGCGGCGGAGGGTGCGGCGCAGGGAGATCTGGGTGGCTCCG

Cre12.g542569
5': ATTGTCAGCCCTAGTCATCAGTGGCTTCGCCACTACCAAGTGTAACTATTGAAGATCCTTTGATCTTTTCTACGGG
3': AGAGCATCGTTTCTTTCCTGTACCAGCCGCCACTGCCCCCGCCGCGGTGGGGAGATCTGGGTGGCTCCG

Cre01.g008550
5': AAAATGCTCAAACGTGTGATTGTGTGTGTATGTTTTTGTGATCGTGTGTCGAAGATCCTTTGATCTTTTCTACGGG
3': AGCAGGCGCTGCGCCACCCCTGGCTGCAGTACCGCTACCGCTCGCCCAAGGGAGATCTGGGTGGCTCCG

Note that the primer sequences for each gene above are all displayed from 5'-3'. The names 5' and 3' refer to positions on the target gene with homology to the 50 bp arms of the respective primers.

#### Supplemental Method 2. BACSearcher

##### A. BACSearcher usage

**Function**

BACSearcher considers all or a subset of Chlamydomonas genes and provides three key pieces of information useful for recombineering a gene of interest using the pipeline detailed in Supplementary Method 1:

(1) The script identifies up to five 50 bp regions chosen from a stretch of sequence 2000-3000 bp upstream of the annotated start codon for each gene. Regions are picked based on low GC content and absence of mono- and dinucleotide repeats. The script reports the reverse complement of each of these regions, any of which can be added to the 5' end of the universal primer GAAGATCCTTTGATCTTTTCTACGGG to produce a target-specific cloning primer. When used in tandem with the 50 bp immediately upstream of the stop codon, prepended to the 5' end of universal primer GGAGATCTGGGTGGCTCCG, these cloning primers can amplify a target-specific recombineering cassette from pLM099 by PCR (see Supplementary Method 1C). Instructions to modify the lengths of these regions are detailed below (see Modifications to the script).

(2) The script reports four pairs of Primer3-generated checking primers for each gene, two that can amplify a short sequence from the 5' end of the gene and two from the 3' end. These can be used to confirm the presence of the 5' and 3' ends of each target gene within a BAC in the Chlamydomonas BAC library.

(3) The script reports the five smallest BACs containing each gene where available. BACs are reported if they cover the region spanning from 3000 bp upstream of the gene to the stop codon. An exception is made in cases where this region extends beyond the ends of a chromosome or scaffold, in which case the region is altered so that it starts/ends at the first/last position of the chromosome/scaffold. These parameters can be modified by the user so that BACs are only reported if, for example, the BAC sequence covers a larger 5' flanking region, or also covers the 3'UTR of the gene of interest. Instructions to make these modifications to the script are detailed below.

Two outputs are generated after each run of the script, one for BAC coverage and one for fosmid coverage of each gene. BAC and fosmid data are taken from the version 5.5 annotation of the Chlamydomonas genome. The fosmid output file contains identical information for (1) and (2), but for (3) the BAC-specific information is replaced by fosmid-specific information. Fosmid plate/well coordinates are not provided. The BAC output for all 17,741 genes in the genome is provided in Supplemental Data Set 1.

**Code availability**

BACSearcher is available as a TXT file in the Supplemental Code ZIP folder. The script and usage instructions are also available in a GitHub repository: https://github.com/TZEmrichMills/Chlamydomonas_recombineering

**Specialist Python modules required**

BACSearcher has been tested using Python 3.6 and requires the following modules to be installed:

| | |
|---|---|
| Gffutils | https://pypi.org/project/gffutils/ |
| Interval tree | https://pypi.org/project/intervaltree/ |
| Biopython | https://pypi.org/project/biopython/ |
| Primer3-py | https://pypi.org/project/primer3-py/ |

**Example usage**

BACSearcher can be initiated from the command line using the following options:

```
./BACSearcher.py \
-p BACs_fosmids.pairs.tsv \
-f Creinhardtii_281_v5.0.fa.gz \
-g Creinhardtii_281_v5.5.gene.gff3.gz \
-l Gene_shortlist.txt \
```

```
-w Bac_wells.txt \
-o Chlamydomonas_BACSearcher_results
```

Where `BACs_fosmids.pairs.tsv` refers to precursor file I (below);
`Creinhardtii_281_v5.0.fa.gz` refers to precursor file II;
`Creinhardtii_281_v5.5.gene.gff3.gz` refers to precursor file III;
`Gene_shortlist.txt` refers to precursor file IV; and
`BAC_wells.txt` refers to precursor file V.
`Chlamydomonas_BACSearcher_results` is the output file name.

**Required precursor files**

I. TSV file containing the coordinates of the start and end of each valid BAC in the library. BACs are included as valid if their start and end sequences are mapped to the same chromosome and are in the correct orientation, (i.e., one end on each strand). This file is provided in the Supplemental Code ZIP folder as BACs_fosmids.pairs.tsv and is also available from the GitHub repository (see Code availability section above).
II. Zipped FASTA file (.fa.gz extension) containing the gene sequences for all Chlamydomonas nuclear genes.
III. Zipped GFF file (.gff3.gz extension) containing version 5.5 annotation information for the Chlamydomonas genome.
IV. (Optional) TXT file containing the Cre IDs for all genes of interest to be processed, one per line, each appended with '.v5.5'. If this file is not provided, BACSearcher will process all nuclear genes and produce a TSV file of the results with the name specified by `-o` (see Example usage, above).
V. TXT file containing the plate and well coordinates of each BAC in the library, in the format 'A-B-C', where A is the plate number, B the row number and C the column number. This file is provided in the Supplemental Code ZIP folder as BAC_wells.txt and is also available from the GitHub repository (see Code availability section).
VI. (Optional) DB file generated from III using the BACSearcher script, which can be used in place of III in future runs.

Genome FASTA and gene annotation GFF files (precursors II and III) are available for download from Phytozome, entitled Creinhardtii_281_v5.0.fa.gz and Creinhardtii_281_v5.5.gene.gff3.gz. The output provided in Supplemental Data Set 1 used precursor files II and III downloaded from Phytozome V12.

When supplied with a GFF file via `-g`, BACSearcher will generate a gffutils database for the GFF file (precursor file VI). BACSearcher can also use this database directly with the `-d` option, saving the effort of regenerating the database.

**Modifications to the script**

The BACSearcher output for all 17,741 Chlamydomonas genes is provided in Supplemental Data Set 1 according to default parameters described in the Function section, above. Users can modify the script to change the length or position of the homology regions that BACSearcher reports, as well as the region that reported BACs should cover for each gene. These modifications are made by adding the additional options `-q`, `-r`, `-s`, `-t`, `-u` and `-v` to the command line before running the script.

(1) Modifying the lengths of the homology arms:
By default, BACSearcher reports 5' and 3' homology regions for each gene that are 50 bp long. To change these lengths to a different value, $x$, the options `-q` and `-r` can be added to the command line:
- `-q` $x$ will change the default lengths of the reported 5' homology regions to an integer, $x$ bp.
- `-r` $x$ will change the default length of the 3' homology region to an integer, $x$ bp.

We recommend using the same values for `-q` and `-r`.

(2) Modifying the size of the upstream native promoter region:
By default, BACSearcher searches for 5' homology regions 2000-3000 bp upstream of the start codon. To change the searched region, the default maximum flank of 3000 can be changed to a different value, $x$, by adding the option, `-s`, to the command line:
- `-s` $x$ will direct the script to search for suitable homology regions in the 1000 bp downstream of an upstream position, $x$ bp. For example, if $x$=5000 the script will search for regions between 4000 and 5000 bp upstream of the start codon of each gene.

If you would like to measure from the start of the 5'UTR instead of from the start codon, the option `-u` can be added to the command line:
- `-u` will direct the script to search for homology regions in the region defined by `-s` but measured from the 5'UTR instead of the start codon. If `-u` is used but `-s` is left undefined, the script will search for homology regions 2000-3000 bp upstream of the 5'UTR.

(3) Modifying the BACs reported for each gene:
By default, the script is set to report BACs that cover the coding region plus the upstream flank for each gene, ignoring the 3'UTR. The upstream flank is defined by `-s` and `-u` (see (2), above) with a default value of 3000 bp upstream from the start codon. If you would like to add a downstream flank of length $x$ bp that any reported BACs should also cover, the option `-t` can be added to the command line:
- `-t` $x$ will direct the script to report only those BACs that cover $x$ bp downstream from the stop codon of each gene.

If you would like to measure the downstream flank from the end of the 3'UTR instead of from the stop codon, the option `-v` can be added to the command line:
- `-v` will direct the script to report BACs that cover a downstream region defined by `-t` but measured from the 3'UTR instead of from the stop codon of each gene. If `-v` is used but `-t` is left undefined, the script will report BACs that cover up to the 3' end of the 3'UTR.

#### B. BAC annotation issue

BACSearcher reports the five smallest BACs per gene that completely cover the gene of interest, plus any additional flanks that the user defines (see Modifications to the script, above). Coverage is implied based on coordinates for each of the two BAC ends; if both ends map onto the same chromosome in the correct orientation (one on each strand), and if one end maps to a position upstream and the other downstream of the region of interest, BACSearcher assumes a continuous BAC sequence exists between these two sites that covers the region of interest. Data for the location of the ends of each BAC are taken from the version 5.5 annotation of the Chlamydomonas genome. These data appear to have been obtained based on the results of a BLAST search, in which a short (<1000 bp) sequence from each BAC end has been queried against the Chlamydomonas genome and the top hits from each search taken as a BAC end location. However, the repetitive nature of the Chlamydomonas genome makes identifying the true ends of each BAC difficult, since a BLAST search based on BAC end sequencing data can produce multiple high-identity hits on the same chromosome. We have identified a handful of cases for which the reported BAC end locations are incorrectly annotated in v5.5 of the Chlamydomonas genome. These may be due either to the wrong BLAST hit having been reported, or due to the query sequence not having been long enough to produce a unique hit site. In support of this observation, we have been able to successfully retrieve a gene (Cre16.g678661) from a BAC (PTQ13193) even though the end coordinates for this BAC do not cover this gene according to the v5.5 annotation. What is more, in this case the BAC ends are recorded in v5.5 as being on the same strand, and so PTQ13193 is not included for consideration by BACSearcher. Our analysis shows that as many as 27.2% of BACs and 36.8% of fosmids are disqualified from consideration by BACSearcher, either because they have end coordinates on different chromosomes or on the same strand of a chromosome, or they only have available coordinates for one end.

Given the potential for misannotation, we recommend two precautions when using this software. (1) If your target gene is not covered by a BAC in the BACSearcher output, we recommend double checking against the archived BAC annotation for version 4 of the Chlamydomonas genome assembly (see the section below for instructions). (2) Prior to commencing the recombineering pipeline, we recommend performing a colony PCR on some or all BAC clones using the Primer3-generated checking primers (see Supplemental Data Set 1) to confirm the presence of your target gene(s) within a BAC.

**Checking BAC coverage using the v4 genome assembly**

(1) BLAST a short sequence from both ends of your target region against the archived version 4 of the Chlamydomonas genome assembly (Chlre4) using the following link: https://mycocosm.jgi.doe.gov/pages/blast-query.jsf?db=Chlre4

(2) Select the top BLAST hit for each sequence to enter the JBrowse view. Below the hit site base positions, find and select the annotation track entitled ‘paired BAC ends’ to view the BACs that cover your query sequence.

(3) If both your query sequences are covered by the same BAC, copy the BAC ID (PTQ number) into the BAC converter to identify well coordinates for this BAC within the library. The converter spreadsheet is available at the following link: https://www.chlamycollection.org/resources/tools/bac-converter/

#### Supplemental Method 3. Bioinformatics software usage

##### A. Tandem Repeats Finder

Tandem Repeats Finder (version 4.09; Benson, 1999) was downloaded from https://tandem.bu.edu/trf/trf.download.html and run from the command line using the following command for all sequences analysed in this work:

```
./trf409.dos64.exe sequence.fa 2 5 7 80 10 20 2000 -h
```

Where `sequence.fa` is a FASTA file containing the genes to be analysed. The first three numbers (`2 5 7`) set the algorithm to be more permissive than recommended in order that repeats with some identity mismatches could be reported. For each dataset analysed, the resulting output was then processed using a custom python script (see Supplemental Method 4A) and analysed using a spreadsheet so that only those repeats with ≥90% identity were incorporated into the results presented in this work. The next two numbers (`80 10`) are the recommended match probability and indel probability settings. Further explanations for all input options can be found on the host website help pages: https://tandem.bu.edu/trf/trf.unix.help.html. The next number (`20`) is the minimum alignment score that a repeat must meet to be reported. This was set to 20 to produce a cut-off such that homopolymer repeats 10 bp long (but no shorter) were included in the output. The final number (`2000`) is the maximum period size detected in base pairs. The `-h` option is provided to generate a data file instead of an html page of the results. Analysis spreadsheets are available upon request.

##### B. Palindrome Analyser

Analysis of inverted repeats was performed using Palindrome Analyser (version 2.6.6; Brázda et al., 2016) available at the following link: http://bioinformatics.ibp.cz:9999/#/en/palindrome. The default settings were modified to report inverted repeats with a maximum mismatch of 1 bp for every 10 bp of stem sequence (equating to a ≥90% identity threshold). Stem sequence lengths were limited to 10-200 bp and loop (spacer) lengths to 0-10 bp. All results were copied into a spreadsheet and analysed to produce values for the average number of repeats per gene or per kilobase, and the percentage of genes containing at least one repeat. The analysis spreadsheet is available upon request.

##### C. WindowMasker

WindowMasker was downloaded as part of the BLAST+ applications from NCBI (version 2.10.0; Morgulis et al., 2006a). These are available as a package from https://ftp.ncbi.nlm.nih.gov/blast/executables/blast+/LATEST/, and additional information for their use is detailed in the user manual accessed using the following link:
http://nebc.nerc.ac.uk/bioinformatics/documentation/blast+/user_manual.pdf.
WindowMasker was run from the command line in two stages using the following options:

```
./windowmasker -mk_counts -in sequence.fa -out sequence.counts

./windowmasker -ustat sequence.counts -in sequence.fa \
-out sequence.intervals.txt -dust true
```

Where `-mk_counts` sets the program to perform a stage 1 read-through of the input sequences (here represented by `sequence.fa`) and outputs a unit counts file, e.g., `sequence.counts`. The `-ustat` command accepts the unit counts file from stage 1 and uses this to mask the input sequence in stage 2. An explanation of the default input options along with full program details can be found at the following link:
https://www.ncbi.nlm.nih.gov/IEB/ToolBox/CPP_DOC/lxr/source/src/app/winmasker/README.
The input settings for all sequences analysed in this work were left as default apart from the `-dust` option; when set to true, this option uses the DUST module (Morgulis et al., 2006b) in addition to the WinMask module. With DUST active, the program masks simple repeats such as tandem and inverted repeats in addition to longer, non-adjacent repeats. By default, the output file (represented here by `sequence.intervals.txt`) contains the name of each sequence in the input file followed by the start and end coordinates for all runs of masked

nucleotides within each sequence. Interval files for all sequences were processed using a custom python script (see Supplemental Method 4B) and analysed in a spreadsheet to produce values for the average number of masked regions per gene or per kilobase, and the percentage of genes containing at least one masked region. Analysis spreadsheets are available upon request.

#### D. Primer3

Primer3 software was used to analyse a dataset of genome-wide ATG-Stop primers in two ways. Firstly, the python module primer3-py (version 0.6.0) was used to produce $\Delta G$ values for predicted secondary structures within and between primers and primer pairs (see Figure 1D, blue bars). Further details on the use of this module can be found here: https://pypi.org/project/primer3-py/. Secondly, the command line version of the Primer3 core (Rozen and Skaletsky, 2000) was used to examine the primary reasons for rejection of those primers in the data set that breached Primer3 thresholds (Figure 1D, orange bar). Primer3 (version 2.5.0) was downloaded as a bioconda package from https://anaconda.org/bioconda/primer3 and run from the anaconda command line using the following commands:

```
./primer3_core –p3_settings_file=path/settings.txt < path/sequences.txt \
> path/output.txt
```

Where `path/sequences.txt` and `path/output.txt` represent pathnames for the input and output files respectively, and `path/settings.txt` is the pathname for a file containing instructions specifying the use of the check_primers module, as well as the parameters this module should use when analysing primer sequences. A list of primers to be analysed was converted to Boulder-IO format using a custom python script prior to analysis (see Supplemental Method 4C). Further details regarding Boulder-IO format and a detailed description of Primer3 and its uses can be found in the manual at the following link: http://primer3.org/manual.html#installMac. Settings used were as default for Primer3Plus (Untergasser et al., 2007) with some minor modifications. Primer3Plus is an enhanced version of Primer3 that acts as an up-to-date online-only interface for the Primer3 core. In order to analyse primers from the command line version of Primer3 using the default Primer3Plus settings, the default Primer3 settings was downloaded as a text file from http://primer3.ut.ee/ and modified to closely align with the Primer3Plus settings (available from http://www.bioinformatics.nl/cgi-bin/primer3plus/primer3plus.cgi, under the general settings tab). Further details on the differences between Primer3 and Primer3Plus can be found in section 5 of the manual at the above link. The resulting settings file was used to analyse all ATG-Stop primers in this work and is shown in full (in Boulder-IO format) below. The key parameter settings to note are as follows:

| Primer3 parameter: | Value: | Notes: |
|---|---|---|
| `PRIMER_THERMODYNAMIC_OLIGO_ALIGNMENT` | 0 | Primer3Plus default. Directs Primer3 to use a scoring-based method for calculating secondary structure alignment scores. |
| `PRIMER_MAX_GC` | 80 | Primer3Plus default. Primers only breach the threshold for GC content if they contain over 80% G/C bases. |
| `PRIMER_TM_FORMULA` | 1 | Directs Primer3 to use the recommended Tm prediction method from SantaLucia (1998). |
| `PRIMER_SALT_CORRECTIONs` | 1 | Directs Primer3 to use the recommended salt correction method from SantaLucia (1998). |
| `PRIMER_MIN_TM` | 40 | Set low to avoid rejecting primers for their predicted melting temperatures. |
| `PRIMER_MAX_TM` | 100 | Set very high to effectively remove any melting temperature restrictions. |

Note that melting temperature restrictions were removed since the dataset of primers we analysed was generated for ATG-Stop cloning of Chlamydomonas gDNA (see Methods). Primers designed against Chlamydomonas gDNA are expected to have higher Tms than recommended based on generic guidelines due to the unusually high GC content of the genome. Removing Tm restrictions enabled a more detailed understanding of secondary structure related issues for these primers.

**Primer3 settings file**

```
Primer3 File - http://primer3.sourceforge.net
P3_FILE_TYPE=settings

PRIMER_FIRST_BASE_INDEX=1
PRIMER_THERMODYNAMIC_OLIGO_ALIGNMENT=0
PRIMER_THERMODYNAMIC_TEMPLATE_ALIGNMENT=0
PRIMER_PICK_LEFT_PRIMER=1
PRIMER_PICK_INTERNAL_OLIGO=0
PRIMER_PICK_RIGHT_PRIMER=1
PRIMER_LIBERAL_BASE=1
PRIMER_LIB_AMBIGUITY_CODES_CONSENSUS=0
PRIMER_LOWERCASE_MASKING=0
PRIMER_PICK_ANYWAY=0
PRIMER_EXPLAIN_FLAG=1
PRIMER_MASK_TEMPLATE=0
PRIMER_TASK=check_primers
PRIMER_MASK_FAILURE_RATE=0.1
PRIMER_MASK_5P_DIRECTION=1
PRIMER_MASK_3P_DIRECTION=0
PRIMER_MIN_QUALITY=0
PRIMER_MIN_END_QUALITY=0
PRIMER_QUALITY_RANGE_MIN=0
PRIMER_QUALITY_RANGE_MAX=100
PRIMER_MIN_SIZE=18
PRIMER_OPT_SIZE=20
PRIMER_MAX_SIZE=27
PRIMER_MIN_TM=40.0
PRIMER_OPT_TM=65
PRIMER_MAX_TM=100
PRIMER_PAIR_MAX_DIFF_TM=100.0
PRIMER_TM_FORMULA=1
PRIMER_PRODUCT_MIN_TM=-1000000.0
PRIMER_PRODUCT_OPT_TM=0.0
PRIMER_PRODUCT_MAX_TM=1000000.0
PRIMER_MIN_GC=30.0
PRIMER_OPT_GC_PERCENT=50.0
PRIMER_MAX_GC=80.0
PRIMER_PRODUCT_SIZE_RANGE=150-250 100-300 301-400 401-500 501-600 601-700 701-850
851-1000
PRIMER_NUM_RETURN=5
PRIMER_MAX_END_STABILITY=9.0
PRIMER_MAX_LIBRARY_MISPRIMING=-1
PRIMER_PAIR_MAX_LIBRARY_MISPRIMING=-1
PRIMER_MAX_SELF_ANY_TH=45.0
PRIMER_MAX_SELF_END_TH=35.0
PRIMER_PAIR_MAX_COMPL_ANY_TH=45.0
PRIMER_PAIR_MAX_COMPL_END_TH=35.0
PRIMER_MAX_HAIRPIN_TH=47.0
PRIMER_MAX_SELF_ANY=8.00
PRIMER_MAX_SELF_END=3.00
```

```
PRIMER_PAIR_MAX_COMPL_ANY=8.00
PRIMER_PAIR_MAX_COMPL_END=3.00
PRIMER_MAX_TEMPLATE_MISPRIMING_TH=-1
PRIMER_PAIR_MAX_TEMPLATE_MISPRIMING_TH=-1
PRIMER_MAX_TEMPLATE_MISPRIMING=-1
PRIMER_PAIR_MAX_TEMPLATE_MISPRIMING=-1
PRIMER_MAX_NS_ACCEPTED=0
PRIMER_MAX_POLY_X=5
PRIMER_INSIDE_PENALTY=-1.0
PRIMER_OUTSIDE_PENALTY=0
PRIMER_GC_CLAMP=0
PRIMER_MAX_END_GC=5
PRIMER_MIN_LEFT_THREE_PRIME_DISTANCE=3
PRIMER_MIN_RIGHT_THREE_PRIME_DISTANCE=3
PRIMER_MIN_5_PRIME_OVERLAP_OF_JUNCTION=7
PRIMER_MIN_3_PRIME_OVERLAP_OF_JUNCTION=4
PRIMER_SALT_MONOVALENT=50.0
PRIMER_SALT_CORRECTIONS=1
PRIMER_SALT_DIVALENT=1.5
PRIMER_DNTP_CONC=0.6
PRIMER_DNA_CONC=50.0
PRIMER_SEQUENCING_SPACING=500
PRIMER_SEQUENCING_INTERVAL=250
PRIMER_SEQUENCING_LEAD=50
PRIMER_SEQUENCING_ACCURACY=20
PRIMER_WT_SIZE_LT=1.0
PRIMER_WT_SIZE_GT=1.0
PRIMER_WT_TM_LT=1.0
PRIMER_WT_TM_GT=1.0
PRIMER_WT_GC_PERCENT_LT=0.0
PRIMER_WT_GC_PERCENT_GT=0.0
PRIMER_WT_SELF_ANY_TH=0.0
PRIMER_WT_SELF_END_TH=0.0
PRIMER_WT_HAIRPIN_TH=0.0
PRIMER_WT_TEMPLATE_MISPRIMING_TH=0.0
PRIMER_WT_SELF_ANY=0.0
PRIMER_WT_SELF_END=0.0
PRIMER_WT_TEMPLATE_MISPRIMING=0.0
PRIMER_WT_NUM_NS=0.0
PRIMER_WT_LIBRARY_MISPRIMING=0.0
PRIMER_WT_SEQ_QUAL=0.0
PRIMER_WT_END_QUAL=0.0
PRIMER_WT_POS_PENALTY=0.0
PRIMER_WT_END_STABILITY=0.0
PRIMER_WT_MASK_FAILURE_RATE=0.0
PRIMER_PAIR_WT_PRODUCT_SIZE_LT=0.0
PRIMER_PAIR_WT_PRODUCT_SIZE_GT=0.0
PRIMER_PAIR_WT_PRODUCT_TM_LT=0.0
PRIMER_PAIR_WT_PRODUCT_TM_GT=0.0
PRIMER_PAIR_WT_COMPL_ANY_TH=0.0
PRIMER_PAIR_WT_COMPL_END_TH=0.0
PRIMER_PAIR_WT_TEMPLATE_MISPRIMING_TH=0.0
PRIMER_PAIR_WT_COMPL_ANY=0.0
PRIMER_PAIR_WT_COMPL_END=0.0
```

```
PRIMER_PAIR_WT_TEMPLATE_MISPRIMING=0.0
PRIMER_PAIR_WT_DIFF_TM=0.0
PRIMER_PAIR_WT_LIBRARY_MISPRIMING=0.0
PRIMER_PAIR_WT_PR_PENALTY=1.0
PRIMER_PAIR_WT_IO_PENALTY=0.0
PRIMER_INTERNAL_MIN_SIZE=18
PRIMER_INTERNAL_OPT_SIZE=20
PRIMER_INTERNAL_MAX_SIZE=27
PRIMER_INTERNAL_MIN_TM=57.0
PRIMER_INTERNAL_OPT_TM=60.0
PRIMER_INTERNAL_MAX_TM=63.0
PRIMER_INTERNAL_MIN_GC=20.0
PRIMER_INTERNAL_OPT_GC_PERCENT=50.0
PRIMER_INTERNAL_MAX_GC=80.0
PRIMER_INTERNAL_MAX_SELF_ANY_TH=47.00
PRIMER_INTERNAL_MAX_SELF_END_TH=47.00
PRIMER_INTERNAL_MAX_HAIRPIN_TH=47.00
PRIMER_INTERNAL_MAX_SELF_ANY=12.00
PRIMER_INTERNAL_MAX_SELF_END=12.00
PRIMER_INTERNAL_MIN_QUALITY=0
PRIMER_INTERNAL_MAX_NS_ACCEPTED=0
PRIMER_INTERNAL_MAX_POLY_X=5
PRIMER_INTERNAL_MAX_LIBRARY_MISHYB=12.00
PRIMER_INTERNAL_SALT_MONOVALENT=50.0
PRIMER_INTERNAL_DNA_CONC=50.0
PRIMER_INTERNAL_SALT_DIVALENT=1.5
PRIMER_INTERNAL_DNTP_CONC=0.0
PRIMER_INTERNAL_WT_SIZE_LT=1.0
PRIMER_INTERNAL_WT_SIZE_GT=1.0
PRIMER_INTERNAL_WT_TM_LT=1.0
PRIMER_INTERNAL_WT_TM_GT=1.0
PRIMER_INTERNAL_WT_GC_PERCENT_LT=0.0
PRIMER_INTERNAL_WT_GC_PERCENT_GT=0.0
PRIMER_INTERNAL_WT_SELF_ANY_TH=0.0
PRIMER_INTERNAL_WT_SELF_END_TH=0.0
PRIMER_INTERNAL_WT_HAIRPIN_TH=0.0
PRIMER_INTERNAL_WT_SELF_ANY=0.0
PRIMER_INTERNAL_WT_SELF_END=0.0
PRIMER_INTERNAL_WT_NUM_NS=0.0
PRIMER_INTERNAL_WT_LIBRARY_MISHYB=0.0
PRIMER_INTERNAL_WT_SEQ_QUAL=0.0
PRIMER_INTERNAL_WT_END_QUAL=0.0
=
```

#### Supplemental Method 4. Bioinformatics python analysis

Scripts are provided as TXT files in the Supplemental Code ZIP folder, and are also available in the associated GitHub repository via the following link: https://github.com/TZEmrichMills/Chlamydomonas_recombineering

##### A. Tandem Repeats Finder output sorter

**Usage**
Tandem Repeats Finder (Benson, 1999) outputs a data file containing lists of repeats found in each sequence supplied to the program. In order to quickly organise the contents of these data files we have written a python script (TRF output sorter, Supplemental Code). TRF output sorter reads the data file produced by Tandem Repeats Finder and reformats the information into a CSV file in which each reported repeat sequence (and its respective analysis information) is present on its own row, preceded by the name of the gene in which it was detected. This enables straightforward analysis of the number, length, type and identity of tandem repeats per gene, as well as the number of genes containing at least one tandem repeat.

##### B. WindowMasker output sorter

**Usage**
WindowMasker (Morgulis et al., 2006a) accepts one or multiple nucleotide sequences and outputs an intervals file containing the name of each entry sequence followed by the start and end positions of each masked repetitive sequence found within the entry (one run per line). We have written a python script, WindowMasker output sorter (Supplemental Code), which uses the python data analysis library (pandas) to simply count the number of intervals per entry sequence, equating to the number of masked repeats per sequence. The script outputs a CSV file to enable the quick generation of values for the average number of masked regions per sequence and number of sequences with at least one masked region.

##### C. Primer3 check_primers output sorter

**Usage**
The Primer3 check_primers module outputs a results summary in Boulder-IO format for each primer or pair of primers supplied in the input. In order to quickly analyse the reasons for rejection for all primers in a dataset we have written a python script, Check_primers output sorter (Supplemental Code). This script extracts the reasons for rejection for all rejected primers and outputs a CSV file, enabling fast quantification of the number and type of any warnings or reasons for rejection produced by the module. The script can be run separately depending on whether the user wishes to analyse pairs of primers against each other or just a list of single primers.
